# Force-Correction Analysis Method for Derivation of Multidimensional Free Energy Landscapes from Adaptively Biased Replica Simulations

**DOI:** 10.1101/2021.02.17.431654

**Authors:** Fabrizio Marinelli, José D. Faraldo-Gómez

**Author notes:** Correspondence should be addressed to: Fabrizio Marinelli or José D. Faraldo-Gómez.

## Abstract

A methodology is proposed for the calculation of multidimensional free-energy landscapes of molecular systems, based on analysis of multiple Molecular Dynamics trajectories wherein adaptive biases have been applied to enhance the sampling of different collective variables. In this approach, which we refer to as Force Correction Analysis Method (FCAM), local averages of the total and biasing forces are evaluated post-hoc, and the latter are subtracted from the former to obtain unbiased estimates of the mean force across collective-variable space. Multidimensional free-energy surfaces and minimum free-energy pathways are then derived from integration of the mean force landscape through kinetic Monte Carlo algorithm. To evaluate the proposed method, a series of numerical tests and comparisons with existing approaches were carried out for small molecules, peptides, and proteins, based on all-atom trajectories generated with standard, concurrent and replica-exchange Metadynamics in collective-variable spaces ranging from one- to six-dimensional. The tests confirm the correctness of the FCAM formulation and demonstrate that calculated mean forces and free energies converge rapidly and accurately, outperforming other methods used to unbias this kind of simulation data.

TOC/Abstract Graphic

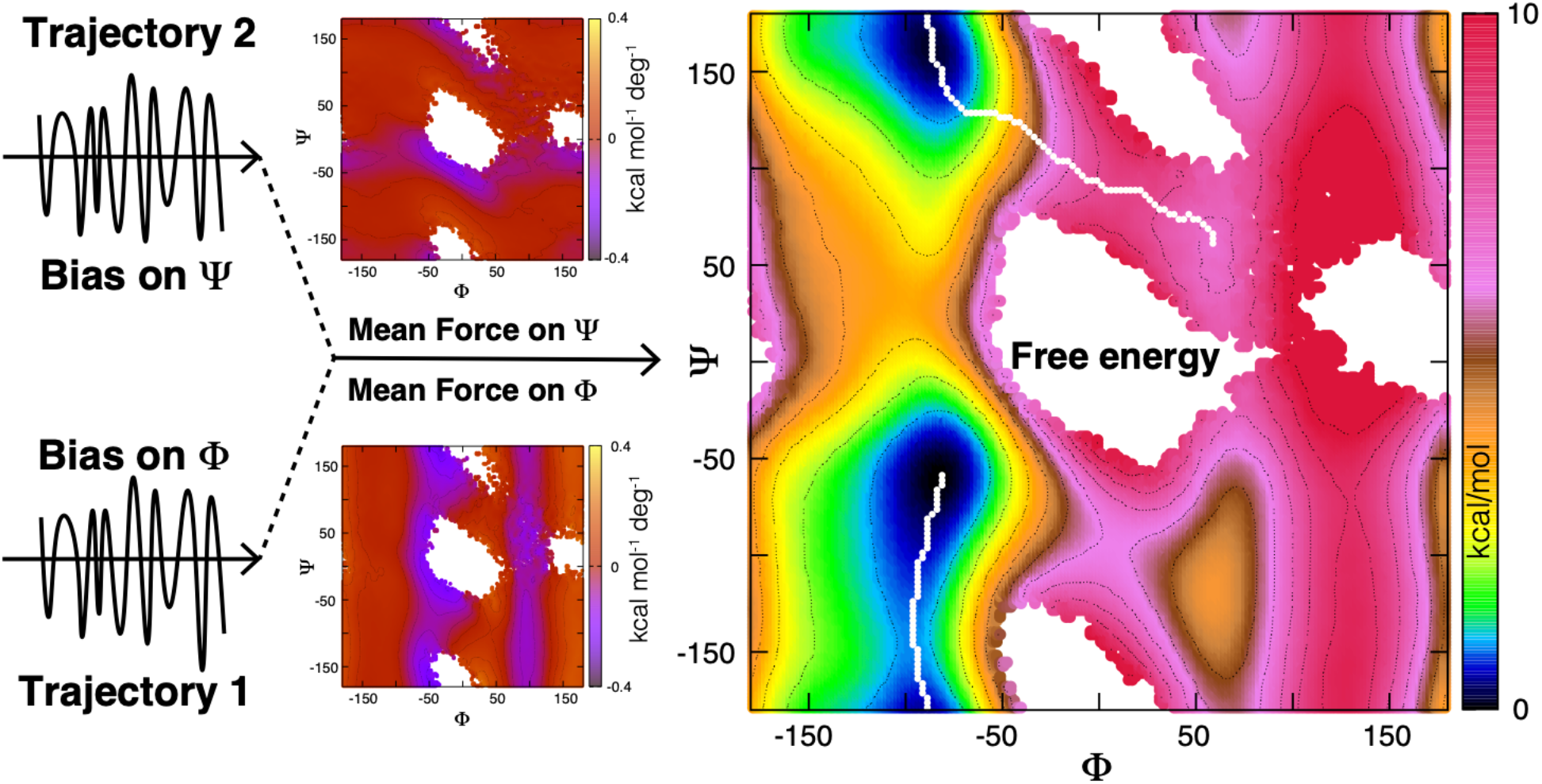

## Introduction

Molecular dynamics (MD) simulations are an increasingly powerful tool to investigate complex chemical and biological mechanisms, and a means to formulate plausible interpretations of experimental measurements that are both physically coherent and grounded on structure at atomic resolution. Too often, however, simulation studies aspire to characterize processes that are too slow for basic algorithms, typically leading to dubious mechanistic inferences based upon largely anecdotal observations. Biased-sampling techniques, by contrast, can yield quantitative information even for slow processes, provided the calculations are designed thoughtfully and analyzed rigorously. This information is ideally in the form of a landscape mapping the free energy of the molecular system as a function of one or more structural descriptors of the process of interest, which are formulated ad-hoc, and are often referred to as collective variables (CVs).^1–4^ The minima and barriers in this landscape intuitively represent the most probable states of the molecular system and the transition pathways that connect them, thus explaining the emergence of a molecular mechanism.

Among a variety of existing biased-sampling techniques, some of the most widely used are umbrella sampling,^5^ adaptive-biasing force (ABF)^6–8^ and Metadynamics.^9–11^ These three methods are alike in that they influence the exploration of collective-variable space by introducing biasing forces in the calculation of atomic trajectories; however, while in US the biasing-potential function is unchanged throughout the simulation, and therefore the resulting forces depend solely on the instantaneous molecular configuration, Metadynamics and ABF are adaptive methods, in that the magnitude and directionality of the bias change gradually as the target CV space is increasingly explored. This adaptability is arguably advantageous, but it also makes a rigorous derivation of free-energies more complex from a theoretical standpoint.^8, 11^ An additional difficulty, common to all biased-sampling techniques, is that it is non-trivial to identify *a priori* what CVs might be the most suitable for the problem at hand; indeed, it is not at all uncommon for intuitive structural descriptors to be entirely ineffective as drivers of configurational sampling. This difficulty has motivated the development of specialized techniques to optimize this choice^12–16^, and to advanced enhanced-sampling protocols that facilitate using a diverse set of tentative CVs, through single^17, 18^ or multiple concurrent biases^19^, applied in individual simulations or through replica-exchange schemes.^20, 21^ Efficient sampling of highly dimensional collective-variable spaces is particularly important in studies of proteins and nucleic acids, as one or two descriptors are often insufficient to adequately to characterize their complex conformational mechanisms.^22, 23^

The challenge ahead is thus to formulate a generic approach to derive multidimensional free-energy landscapes from an arbitrary number of MD trajectories computed with different adaptive-biasing schemes and defined for multiple CVs. In previous applications, we and others have tackled this problem using the Weighted Histogram Analysis Method (WHAM)^24^, an approach that had been (and continues to be) extensively applied in the context of non-adaptive biasing methods (such as umbrella sampling). The underlying concept is that the effect of the biasing potentials can be removed in post-processing by rebalancing the statistical weights assigned to each of the configurations sampled in the trajectories, in reflection of the ‘effective’ bias that was required to attain those configurations; the corrected unbiased sampling can be then combined to obtain the best estimate of the free energy. For adaptive techniques such as Metadynamics, however, the application of WHAM entails a somewhat arbitrary definition of what “effective” bias is, which depends on the specific scheme adopted. For example, in standard Metadynamics this effective bias has been considered to be the time-average of the biasing potential after a certain equilibration time,^11, 25, 26^ while in well-tempered Metadynamics it has been considered to be equal to the bias potential at the end of the simulation.^11, 27^ Although alternative formulations of WHAM that circumvent the definition of an effective bias are conceivable, they are likely to result in significant numerical errors, as they would require iterative determination of large number of shift constants. For similar reasons, related analysis tools such as dynamic histogram^28, 29^ or transition-based reweighting ^30, 31^ have also not been straightforward to apply in the context of adaptive biases.

Building upon the umbrella-integration method^32^ and variants thereof formulated for analysis of adaptively-biased simulations^33–36^, we propose an alternative general approach for the calculation of multidimensional free energy landscapes, based on the calculation of unbiased estimates of the free-energy gradient or mean force. This approach, which we refer to as Force Correction Analysis Method (FCAM), does not require that an effective potential be defined, and is numerically stable. To demonstrate the validity and performance of this methodology, we compare it with existing analysis methods for simulation data obtained for multiple systems and different adaptive sampling schemes of increasing complexity. The cases considered range from single trajectories of simple molecular systems, enhanced to explore one or two CVs, to complex replica-exchange schemes designed for multi-dimensional characterization of polypeptides and proteins.

### Theory

#### Basic concepts

To introduce the formulation, we assume to have a set CVs, 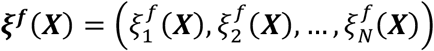, that are functions of the molecular configurations, ***X***. Let us define the free energy as a function of those CVs as:^37, 38^

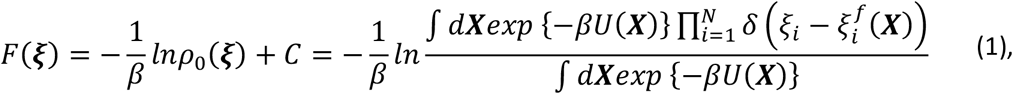

where *ρ*_0_(***ξ***) is the unbiased probability density in the space of the CVs (*C* is a constant), *U*(***X***) is the simulation potential-energy function and *β* = 1/*k*_*B*_*T*, in which *k*_*B*_ is the Boltzmann constant and *T* the temperature. The mean force is the negative of the free-energy gradient:

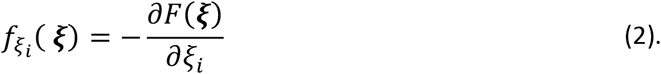

When derived from unbiased molecular dynamics (MD) trajectories, probabilities and free energies are calculated by approximating the Dirac delta function in Eq. 1 with a simple rectangular function or with a density kernel, *K*_*h*_ (***ξ*** – ***ξ***^***f***^(***X***)), e.g. a Gaussian function. This kind of density estimator results in smooth free energies and mean forces, whose accuracy in capturing the correct value depends on the kernel shape and bandwidth. Integrating over degrees of freedom orthogonal to the CVs and using convolution rules, the smoothed mean force can be expressed as:

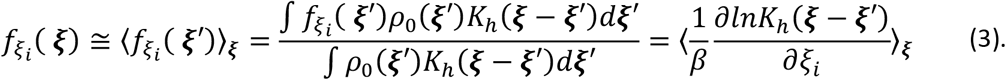

The notation ⟨…⟩_***ξ***_ stands for local average around ***ξ***, which can be calculated as an average over the configurations sampled in the MD trajectory, each weighted proportionally to the density kernel, *K*_*h*_ (***ξ*** – ***ξ***^***f***^(***X***_*t*_)), where ***X***_*t*_ denotes the atomic coordinates at time *t*. The effect of the density kernel is to suppress the weight given to configurations that deviate from ***ξ***. For a Gaussian function of width *σ*, Eq. 3 becomes 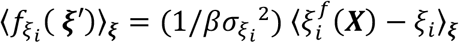. Note that this configurational average is the same as that resulting from an alternative simulation wherein a biasing potential in the form of a harmonic restraint 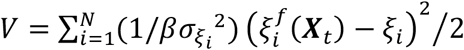 is used to confine the trajectory around ***ξ***. That is, in a restrained ensemble the biasing forces counterbalance on average the intrinsic mean forces, and therefore analysis of the former provide an estimate of the latter.^39, 40^ In the following we will elaborate on this relationship as it applies to adaptively biased simulations. The goal of the derivation is the generalization of Eq. 3 to account for the presence of time-dependent biasing forces applied to the simulation, under a near-equilibrium assumption, and on how to the combine different estimates arising from multiple trajectories to obtain the optimal estimate of the mean force.

#### Mean force estimates from multiple simulations with time-dependent biases

Let us consider the case of a set of MD trajectories, each denoted with the index *r*, wherein a time-dependent biasing potential, *V*_*r*_(***ξ***, *t*), is applied, which may entail different schemes (e.g. standard or well-tempered Metadynamics^11, 41^) or act on different subsets of the CVs considered (as in bias-exchange Metaynamics^20^). Additionally, as expected for properly set Metadynamics^9–11^ or ABF simulations^6–8^, we assume a slow time variation of *V*_*r*_(***ξ***, *t*) such that the simulations remain constantly near equilibrium.

In our previous studies,^25, 42–45^ we used WHAM^24^ to unbias the configurational sampling obtained in these types of simulations and thereby calculate the free energy as a function of the CVs. To do so, the unbiased probability density, 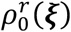, for trajectory *r*, was derived according to umbrella sampling re-weighting^5, 24^ based on an effective bias potential, 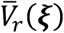, i.e. by adjusting the statistical weight of each configuration sampled. That is:

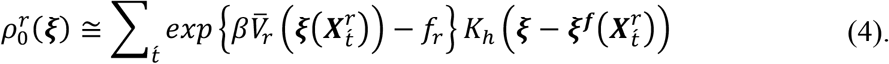

in which 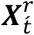 is the configuration of trajectory *r* at time 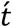 and ƒ_*r*_ is a shift constant evaluated iteratively. The density kernel 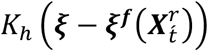 was a simple rectangular function, i.e. 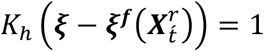 if 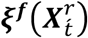 belongs to a bin centered in ***ξ***, and zero otherwise. The free energies were then determined by combining the unbiased probabilities from all trajectories:

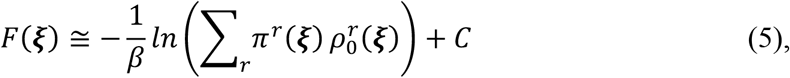

in which the terms *π*^*r*^/(***ξ***) ensure a minimal error on the free energy (according to Poisson’s statistics).

Even though WHAM is in principle a rigorous framework, it is not quite adequate for time-dependent biases in that there is no unique definition of the effective potential 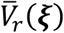 across different adaptive biasing schemes. For example, for standard Metadynamics it has been considered that 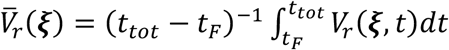, where *t*_*tot*_ is the total simulation time and *t*_*F*_ (filling time) is an equilibration time after which the biasing potential is approximately stationary.^11, 25, 26^ However, for well-tempered Metadynamics, the assumption is that 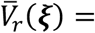 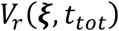 instead.^11, 27, 45, 46^ A possible strategy to circumvent this somewhat arbitrary defintion would be to use the instantaneous bias potential, *V*_*r*_(***ξ***, *t*), and apply Eq. 4 for small simulation intervals, Δ*t*, e.g. at any bias potential update. However, in practice this would imply a very large number of additive constants *f*_*r*_ (one set for each update) that would have to be iteratively determined, which is hardly feasible numerically. Hereafter, we report the basic derivation of the FCAM approach, which does not require the definition of an effective potential nor the iterative evaluation of additive constants.

Let us apply the analogous expression of Eq. 1, through the density kernel *K*_*h*_ (***ξ*** – ***ξ***^***f***^(***X***)), on a time interval, Δ*t*, of trajectory *r* that is short enough so that the time dependence of the bias potential can be neglected (e.g. Δ*t* is the pace of the bias potential time-update). Owing to the presence of *V*_*r*_(***ξ***, *t*), this operation will generally produce a biased estimate of the free energy:

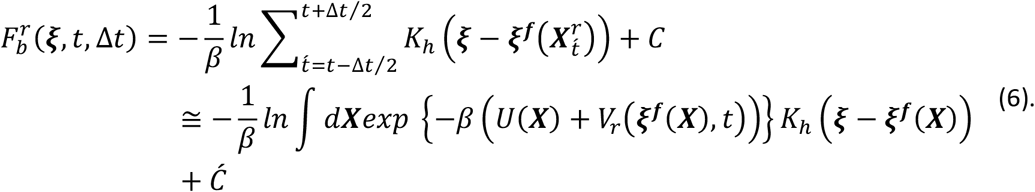

In which the right-hand side of Eq. 6 arises from the equilibrium assumption^5, 32^ (and *C*, *Ć* are constants). To relate Eq. 6 to the unbiased free energy is convenient to integrate over the degrees of freedom orthogonal to the CVs: 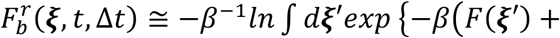 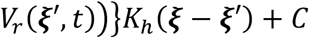. Taking the derivatives of the latter expression (and using convolution rules), we obtain that the unbiased mean force, 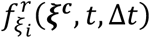, can be recovered from the biased mean force by simply subtracting the local average of the biasing forces:

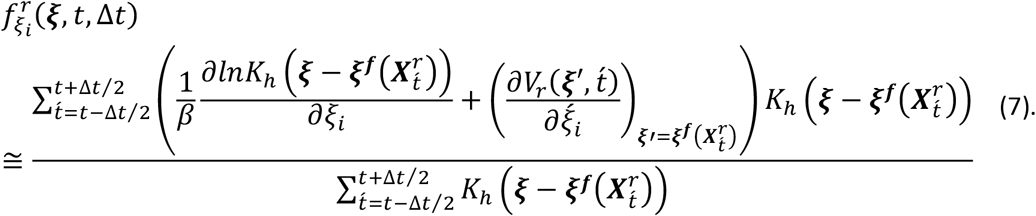

A general expression of the mean force can now be obtained as the weighted average^32^ of all the unbiased estimates, 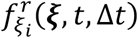, obtained for all time intervals and simulations trajectories, according to the weights 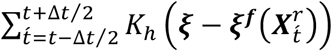:

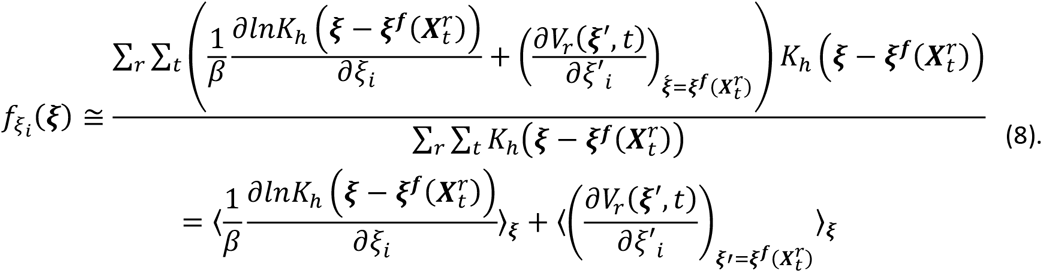

The general meaning of Eq. 8 is that the mean force is the result of the total force inferred from the combined probability density of all simulation trajectories, corrected by subtracting the local average across all trajectories, ⟨…⟩_*ξ*_, of the instantaneous forces originating from the different biasing potentials applied. Note that in contrast to WHAM (Eq. 4), Eq. 8 does not entail effective biasing potentials nor additive constants that must be iteratively evaluated. Hence, it provides a more general formulation valid for any type and combination of adaptively biased simulations, provided they fulfill the near-equilibrium assumption. For Metadynamics and ABF, near-equilibrium conditions are fulfilled when the biasing potential is updated slowly and even more so, after the simulation has explored the relevant regions of the CVs space, so that the biasing forces oscillate around well-defined values.

As in Eq. 3, the effect of the density kernel is to produce a smoothing of the mean forces, hence, a specific selection and tuning of the latter kernel may become particularly advantageous for accurate high dimensional analysis.^47^ Another possibility for large dimensionality is to apply a deconvolution scheme (e.g. based on Fourier analysis) to infer the correct value of the mean force. In all the applications presented in this work, mean forces were calculated according to a simple density kernel selected as a product of Gaussians (with standard deviations 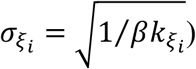 for which Eq. 8 becomes:

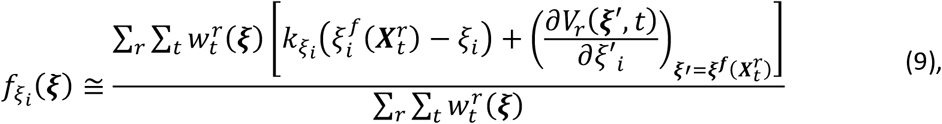

where 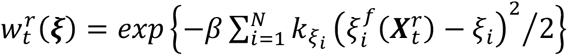, in which the terms 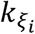 can be considered as smoothing parameters that must be selected large enough so as to capture the relevant features of the underlying free energy. The previous equation (Eq. 9) resembles the expression introduced by Marinova & Salvalaglio to calculate the mean forces from Metadynamics simulations,^36^ where, in contrast to Eq. 9, the biasing force term was estimated as 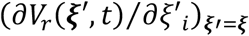. These two approximations become equivalent in the limit of small bandwidth of the density kernel. However, using the instantaneous value of the biasing force (as in Eqs. 8-9), 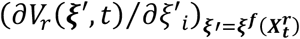, is in our view more computationally convenient as it is often made directly available by enhanced-sampling simulation programs.^48–52^

As the mean force given by Eq. 8-9 arises from a local average, error estimates can be achieved through block averages or autocorrelation analysis^8^. Assuming the same autocorrelation time for all trajectories and time intervals, the statistical error would be proportional to 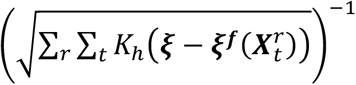, where the sum of the weights is associated to the number of effective frames around ***ξ***.

It is worth pointing out that mean forces can be alternatively evaluated using the formulation adopted in ABF, namely as a local average, around ***ξ***, of the instantaneous forces arising from the simulation energy function, *U*(***X***), and effectively projected along the CVs.^6–8, 53–55^ Assuming equilibrium condition, such local average would not be affected by any bias potential that depends only on the CVs,^51^ hence it could be in principle applied in the context multiple trajectories and biases. This notwithstanding, implementation difficulties and/or inapplicability in presence of constraints and multiple related CVs,^8^ has made this analysis unavailable for several types of CVs and simulation programs. Here, we use estimates based on these mean instantaneous forces (MIF) to evaluate the accuracy and convergence rate of estimates obtained with FCAM for simple mono and two-dimensional examples (see applications to butyramide and deca-L-alanine). In particular, we used the implementation available in NAMD^56, 57^ through the Colvars module,^51^ from which mean instantaneous forces are derived as:^54, 55^

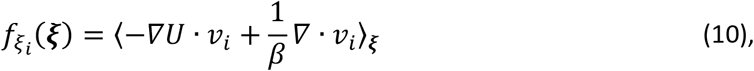

in which the term −*∇U* denotes the atomic forces and *v*_*i*_ is a vector field satisfying a set of orthonormality/orthogonality conditions: *v*_*i*_ · *∇ξ*_*j*_ = *δ*_*ij*_ and *v*_*i*_ · *∇σ*_*k*_ = 0, where *σ*_*k*_(***X***) = 0 defines a constraint applied to the simulation.

#### Multidimensional free energy landscapes from mean forces

After evaluating the mean forces on a dense set of CVs configurations 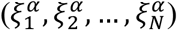, e.g. placed on a regular grid, the free energy can be calculated according to Eq. 2 by integration. To do this accurately and efficiently across multiple dimensions, we used a kinetic Monte Carlo (KMC) scheme that resembles the approach introduced by Hénin et al.^55^. The KMC simulations are based on a transition rate between neighboring bins, *α* and *β*, that satisfies the detailed balance^25^:

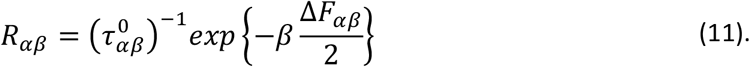

Assuming the transitions dynamics between bins is well approximated by the Markovian model described by Eq. 11, the latter can be also used to obtain kinetic information, provided 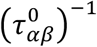 reflects the diffusion rate.^25^ The main purpose of this work, however, is to obtain accurate evaluations of the free energy, which is invariant under changes of 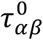; hence, we arbitrarily set the pre-exponential term as a geometric factor: 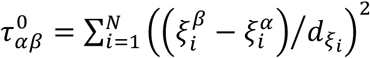, where 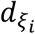 are the bin widths of the grid. The free-energy difference between *α* and *β*, Δ*F*_*αβ*_ = *F*(***ξ***^***β***^) – *F*(***ξ***^***α***^), is obtained from the mean forces using finite differences:

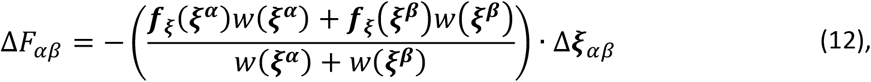

where 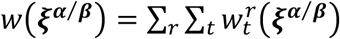 are the total weights of each of the two bins (see Eq. 9) and 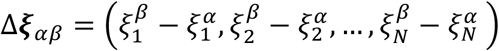 is the inter-bin displacement vector in the space of the CVs.

Finally, free energies can be estimated as in Eq. 1 using the bin probabilities calculated over a long KMC trajectory, 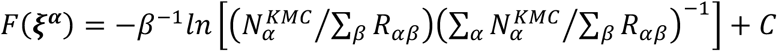, where 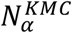 is the number of times the KMC trajectory visits bin *α*. The advantage of using KMC simulations is that, once neighboring bins are assigned, the interconnected regions of the CVs space are automatically identified during the KMC trajectory. Furthermore, statistical convergence can be easily achieved as running long KMC simulations (for billions of steps) is not computationally demanding. It is worth mentioning that alternative approaches can be also used for accurate and efficient derivation of the free energy from the mean forces, based for example on radial-basis function fitting^40^ or on the Poisson equation formalism (the latter is particularly suited for 2-3 dimensions)^58^.

#### Ensemble averages and free energy as function of alternative CVs

Once the free energy as a function of the CVs included in the biasing scheme has been derived (e.g. with FCAM), the free energy as a function of alternative CVs, as well as unbiased ensemble averages of generic observables, can be calculated considering that the sampling distribution remains unbiased within a small bin of the original CVs. Hence, the ensemble average of an observable O can be calculated as^25^:

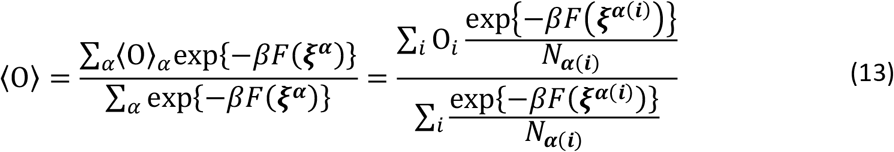

where *F*(***ξ***^***α***^) is the free energy as a function of the original CVs evaluated in a bin *α* and ⟨O⟩_*α*_ is the mean value of O in the same bin. The right-hand side of eq. 13 shows that the latter average is equivalent to a weighted average over the simulation frames of all trajectories with weights provided by exp{−*βF*(***ξ***^***α***(***i***)^)}/*N*_***α***(***i***)_}, where *α*(*i*) is the bin assigned to frame *i* and *N*_***α***(***i***)_ is the cumulative number of frames in bin *α*. Similarly, the free energy as a function of alternative CVs, *F*(***η***), can be derived from the histogram along ***η*** evaluated according to the previous weights:

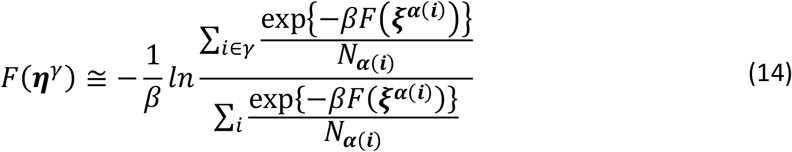

#### Minimum free energy paths calculation across multidimensional landscapes

The minimum free-energy path between two states in the CVs space, *α*_1_, *α*_*N*_, is the most probable mechanism of interconversion between those states.^37^ Assuming this mechanism arises from a series of stochastic transitions between bins in CVs space, the minimum free energy path is that maximizing the time-independent probability of connecting the two states.^37, 59^ This probability is given by the product of the normalized pairwise transition probabilities between bins, which can be derived according to the KMC rates reported in Eq. 11 ^43, 59^:

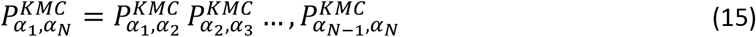

in which 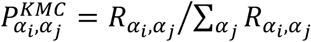 is the probability to observe a transition between *α*_*i*_ and *α*_*j*_. The minimum free-energy pathways reported in this work were obtained using first a global search with KMC trajectories and then a refinement step in which new pathways are generated by randomly selecting two intermediate configurations along the current pathway and running a KMC trajectory that connects them. The sampled pathways are accepted or rejected according to a Monte Carlo (MC) scheme that samples the distribution 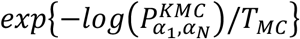, in which *T*_*MC*_ represents a dimensionless temperature factor. Different MC runs are carried out by gradually reducing *T*_*MC*_ using a simulated annealing protocol to converge to the most probable pathway.

### Computational details

Metadynamics simulations of butyramide and deca-L-alanine were carried out at 298 K with NAMD versions 2.12 to 2.14^56, 57^ and colvars,^51^ using the CHARMM27/CMAP force field.^60, 61^ Butyramide was simulated in solution, in a periodic cubic box with 1,467 water molecules, at constant pressure (1 bar), using a 1-fs integration timestep. The simulations of deca-L-alanine were carried out in the gas phase, with a 0.5-fs timestep^7^. To permit the correct evaluation of instantaneous forces according to Eq. 10, no constraints were used on any of the chemical bonds in either butyramide or deca-L-alanine ^8^. Short-range electrostatic and van der Waals interactions were calculated with Coulomb and Lennard-Jones potentials, respectively, cut off at 12 Å. For butyramide, long-range electrostatic interactions were calculated with the PME method. In this system, the bias potentials were constructed by adding Gaussians of widths 5° and 10° for standard and concurrent Metadynamics, respectively. In the case of deca-L-alanine, the Gaussians width was set to 0.5 Å. Gaussians were added every 1 ps and 0.5 ps for butyramide and deca-L-alanine, respectively. Multiple sets of simulations were carried out at different values of the Gaussian height, ranging from 0.025 to 5 kcal/mol for the butyramide simulation system and 0.005 to 0.5 kcal/mol for deca-L-alanine (**Fig. 3** and **Fig. 4**). For the latter, reflective conditions of the bias potential and confinement quadratic restraints were applied at the boundaries (11 and 32 Å) to ensure that stationary conditions are reached^62^. The Bias-Exchange Metadynamics method^20^ was used to simulate the alanine dipeptide and the Ace-Ala_3_-Nme peptide in solution. These simulations were carried out with GROMACS 4.5.5/PLUMED,^48, 50, 63^ and with a timestep of 2 fs. Short-range electrostatic and van der Waals interactions were cut off at 9 Å; long-range electrostatic interactions were calculated with PME. In both cases the bias potentials were constructed using Gaussians of height 0.024 kcal/mol and width 5.7°, added every 2 ps. Exchanges between replicas were attempted every 2 ps. The alanine dipeptide simulation used the CHARMM27 force field,^60^ and includes 877 water molecules while for the Ace-Ala_3_-Nme peptide we used the AMBER03 force field^64^ and 1052 water molecules; both systems are enclosed in a periodic cubic box, and were simulated at 298 K and 1 bar. Computational details for the simulations of the SH3-SH2 domain-tandem of the Abl tyrosine kinase have been reported elsewhere.^44^ Statistics for the calculation of mean forces was collected every simulation step for simulations of butyramide and deca-L-alanine, every 100 steps for alanine dipeptide and Ace-Ala_3_-Nme peptide, and every 250 steps for the SH3-SH2 tandem.

## Results and Discussion

### One-dimensional test cases demonstrate FCAM is highly accurate

To evaluate the numerical accuracy of the proposed approach, we considered two one-dimensional test cases. We first examined the isomerization of butyramide in water, because it is a simple molecular process for which mean forces and free energies can be obtained readily with existing ‘gold-standard’ methods, such as umbrella sampling or long-time conventional MD. To that end, we first carried out 10 independent Metadynamics simulations (30 ns each) using the Φ dihedral angle as the enhanced CV (**Fig. 1**). That is, a time-dependent bias potential was gradually constructed as a sum of Gaussians centered on the values of Φ visited as each of the trajectories progresses^9–11^. After a certain equilibration period, referred to as ‘filling time’ (~6 ns in this case), the bias potential becomes stationary and oscillates around the negative of the free energy profile along Φ, which leads to approximately uniform sampling of Φ^9–11^.

**Figure 1.**
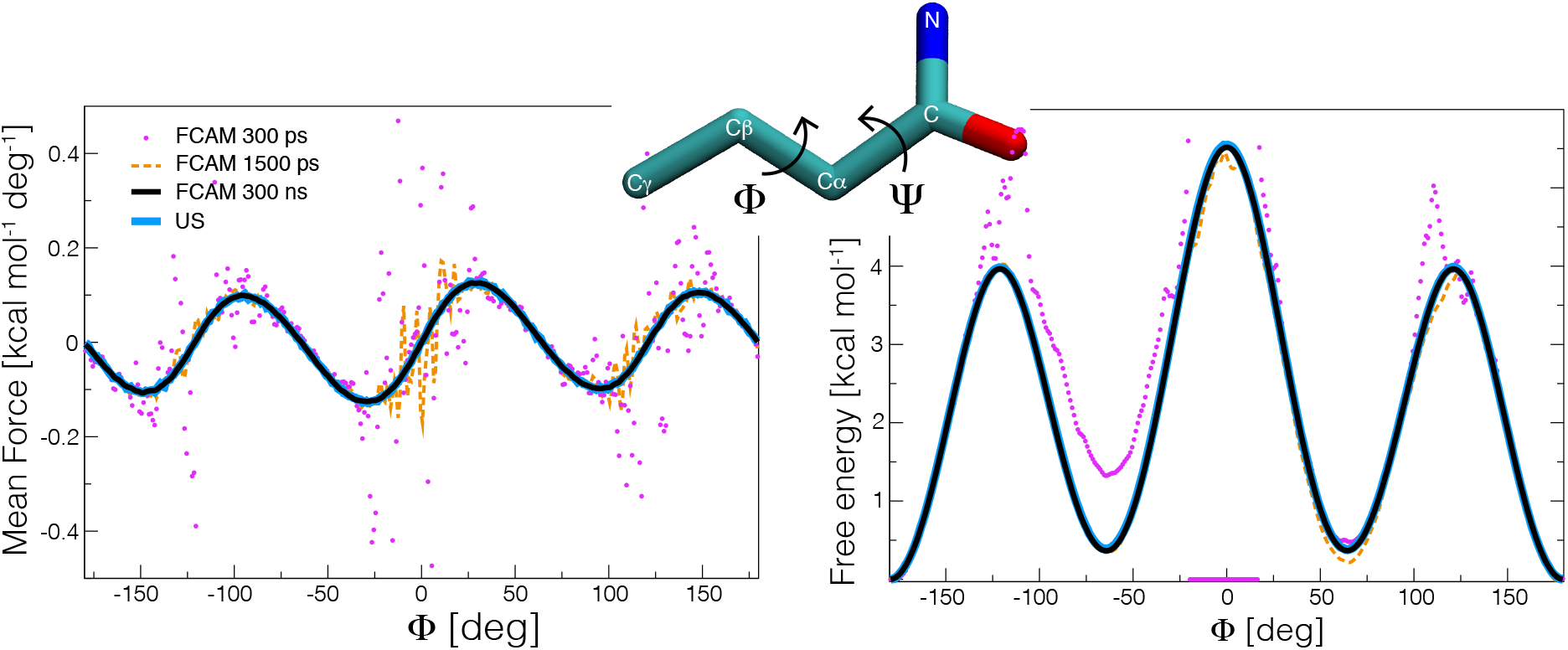
Evaluation of the Force Correction Analysis Method (FCAM), for Metadynamics simulations of solvated butyramide. The butyramide molecule and its Φ and Ψ dihedral angles are depicted in the inset. The Metadynamics biasing potential was constructed using Gaussians of height 0.025 kcal/mol along the Φ dihedral angle (Methods). From left to right, the figure shows mean-force and free-energy profiles (in increments of 1°) calculated with FCAM (*magenta*, *orange*, *black*), using different sampling-time intervals (Eq. 9, with a Gaussian standard deviation of 1° for the density estimator). The FCAM estimates are contrasted with analogous data obtained with a 140-ns umbrella sampling (US) simulation^5^ (*blue*), wherein a static biasing potential, V_US_(Φ), was continuously applied; the specific V_US_(Φ) function used is equal to the biasing potential constructed with a 20-ns Metadynamics simulation. The free energy in this case was estimated using standard histogram reweighting^5^: F_US_(Φ) = −β^−1^ ln (ρ_US_(Φ) exp {β V_US_(Φ)}), where ρ_US_(Φ) is the biased histogram along Φ obtained from the 140-ns US trajectory (using a binning width of 1°). The associated mean forces were obtained by numerical differentiation from F_US_(Φ).

We then used the FCAM method to process this trajectory data and obtain an unbiased estimate of the mean force as a function of the value of Φ, from which the free-energy profile along Φ was derived by integration. As mentioned, we also calculated both mean-force and free-energy profiles with umbrella sampling (US). Comparison of the results demonstrates that the estimates derived from FCAM analysis of the Metadynamics trajectories perfectly match those obtained with US (**Fig. 1**). The mean discrepancy between these methods is comparable to the statistical error of the US estimates, which is 0.002 kcal mol^−1^ deg^−1^ and 0.01 kcal mol^−1^ for mean force and free energy, respectively (based on block analysis). It is also reassuring that for this simple system estimates derived with FCAM converge after a few ns, which is considerably faster than the abovementioned ‘filling time’ (**Fig. 1**). It is worth noting that once an accurate estimate of the free energy along a given CV has been obtained, the free energy along a different CV can be recovered using a reweighting scheme (Eq. 14), provided the latter CV is also adequately sampled in the trajectory. For example, in this case this reweighting procedure can be used to obtain the free energy profile along the Ψ dihedral angle (**Fig. 1**), even though sampling of this angle was not directly enhanced in the original Metadynamics trajectories (**Fig. S1**).

To verify that FCAM is similarly accurate on a slightly more complex system and a different type of CV, we assessed the free energy of reversible unfolding of a deca-L-alanine helix, which has been utilized as a benchmark test case for free-energy simulation methods.^7, 8, 55, 65^ As for butyramide, we first carried out 10 independent Metadynamics simulations (of 100 ns each) biasing in this case the end-to-end distance of the peptide, so as to sample reversibly the transition between α-helical and extended conformations (i.e., distances smaller or larger than 16 Å). We also carried out an independent US calculation, for comparison.

From the data shown in **Fig. 2**, it is clear that also for this system mean forces and free energies calculated with FCAM match the US estimates (within the statistical error of 0.05 kcal/mol^−1^ Å^−1^ and 0.1 kcal/mol, respectively), and closely resemble results reported elsewhere.^7, 8, 55, 65^ In contrast to butyramide, however, the sampling times required for the FCAM estimates to converge are slightly longer than the filling time of the Metadynamics bias (~20 ns); this observation simply reflects the much larger free-energy difference between the endpoints of this conformational transition (**Fig. 2**).

**Figure 2.**
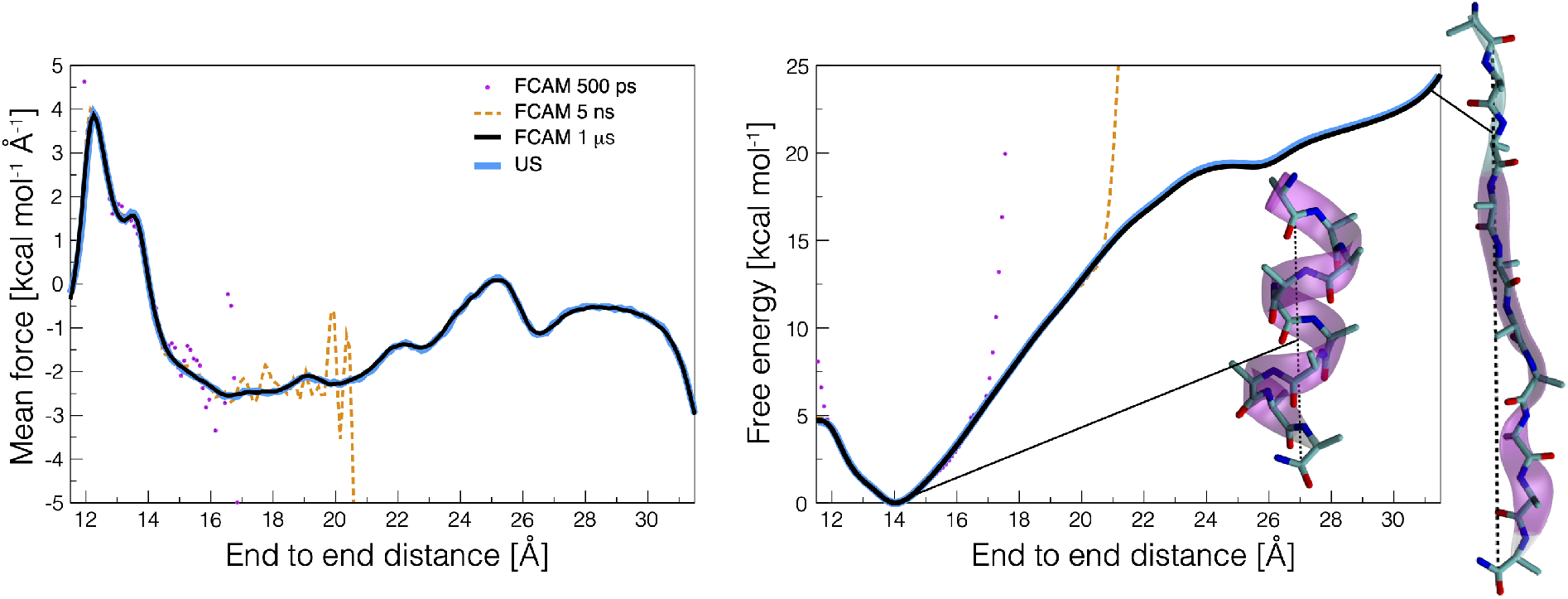
Assessment of FCAM for Metadynamics simulations of a deca-L-alanine peptide enhanced to sample the end-to-end distance (i.e., the distance between the carbon atoms of the carbonyl groups of residues 1 and 10). Molecular configurations of the helical and extended states are depicted in the insets. From left to right, the figure shows mean-force and free-energy profiles calculated with FCAM (Eq.9, with a Gaussian standard deviation of 0.1 Å for the density estimator), using different sampling-time intervals. The FCAM estimates (*magenta*, orange, *black*) are contrasted with analogous data (*blue*) obtained with 10 independent US simulations, 100 ns each. The static biasing potential applied in these US simulations was constructed with 10 Metadynamics simulations (60 ns each), as previously described.^66^

### FCAM converges rapidly and accurately in near-equilibrium condition

An alternative approach for calculating mean-force and free-energy profiles from adaptively-biased MD trajectories is to directly evaluate the effective projection of the mean instantaneous forces (MIF) acting in the molecular system on the space of the CVs of interest.^7^ This approach is limited to simple CVs such as dihedral angles and distances (and subject to other restrictions, e.g., CVs must be orthogonal to each other and any other constraints in the system); however, in such cases this projection can be defined analytically (Eq. 10) and hence these MIF estimates can be thought as ideal. Therefore, a way to evaluate the FCAM method (which is not subject to those restrictions) is to make a direct comparison with MIF estimates for systems amenable to both methodologies, such as the two cases studied in the previous section.

Because FCAM is meant to be applied in the context of adaptively-biased sampling, a key aspect that is important to evaluate is how the time-fluctuations in the amplitude of the applied biasing potential impact the convergence and accuracy of the mean-force estimates. In Metadynamics simulations, these time-fluctuations can be regulated by adjusting the height (*w*) of the Gaussian functions used to construct the biasing potential. This parameter effectively controls the degree to which the system is permitted to deviate from equilibrium, as it determines the magnitude of the local biasing forces added to the simulation per unit of time; indeed, it can be shown that the standard deviation of the time-fluctuations of the biasing potential, σ_b_, increases proportionally with 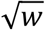.^67, 68^

To evaluate this question for the butyramide test, we generated and analyzed a series of Metadynamics trajectories constructed with Gaussian heights ranging from 0.025 to 5 kcal/mol, analogous to those discussed in the previous section; the corresponding σ_b_ values thus range from 0.1 to 4.6 kcal/mol (**Fig. S2**). In this range, mean-force and free-energy profiles estimated with FCAM match very well the MIF estimates, as well as the ‘gold-standard’ US calculations reported above (**Fig. 3AB**). Analysis of the sampling-time dependence of the errors in the FCAM and MIF estimates with respect to the US estimates confirms both approaches have comparable convergence rate and accuracy (**Fig. 3DE**). In contrast, free-energy profiles directly inferred from the time-average of the Metadynamics biasing potential, while in reasonable agreement the other methods (**Fig. 3C**), are overall less accurate (brown lines in **Fig. 3F**), even after the filling time.

**Figure 3.**
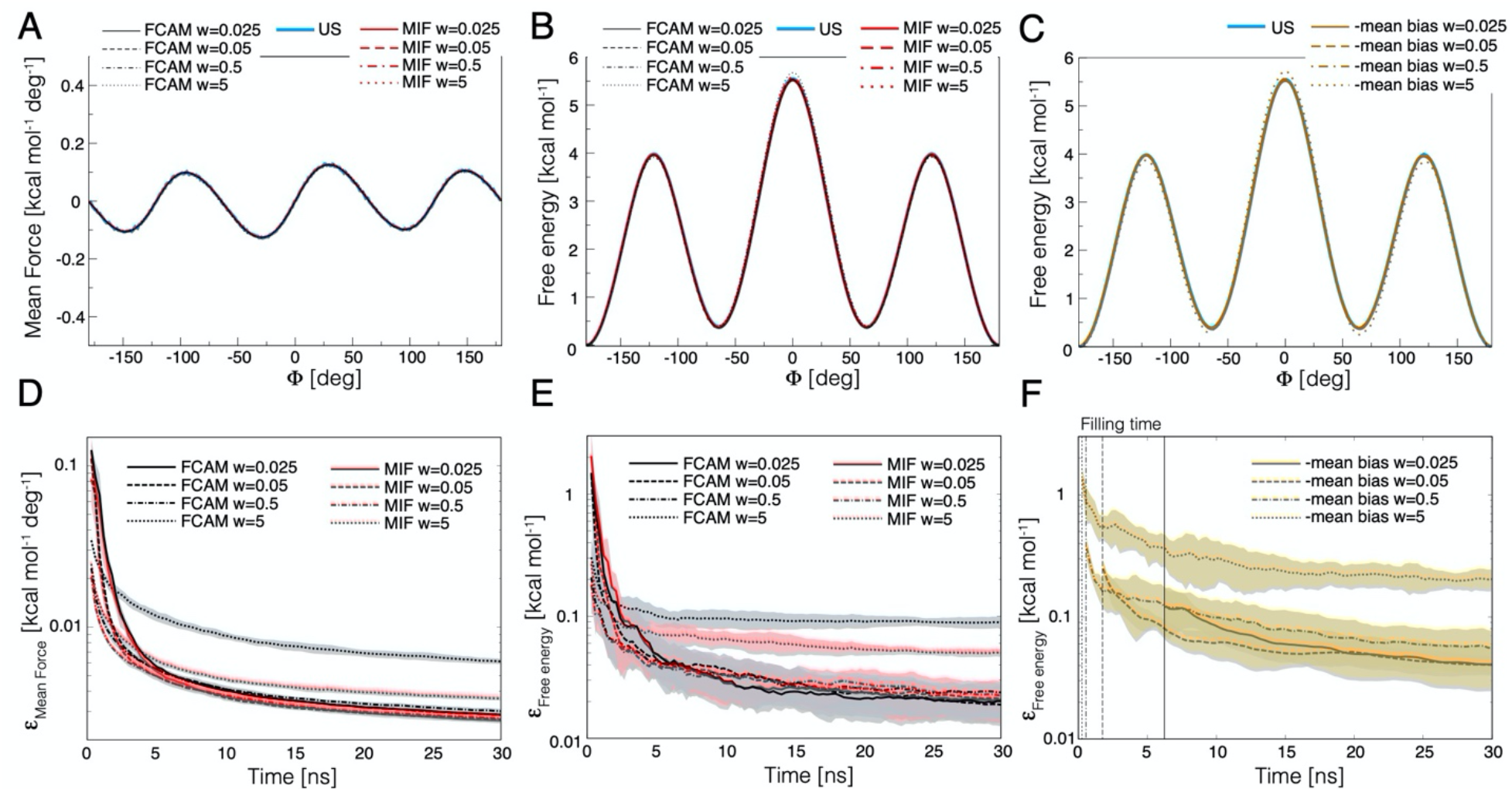
Comparison of FCAM and MIF convergence rates and accuracy, for Metadynamics simulations of solvated butyramide generated with different amplitudes of the time-fluctuations of the biasing potential acting on the Φ angle. (**A**) Mean-force profiles and (**B, C**) free-energy profiles estimated with FCAM (Eq. 9, *black/gray*), with MIF (Eq. 10, *red*) and from the time-average of the Metadynamics biasing potential after the filling time (*brown*),^11, 25, 26^ for different values of the height of the Gaussians used to construct that biasing potential (*w*). Each profile was calculated from the combined sampling of 10 independent Metadynamics trajectories of 30 ns each. Analogous profiles obtained with an umbrella-sampling (US) calculation (same as Fig. 1) are shown for comparison (*blue*). (**D, E, F**) Sampling-time dependence of the error in the FCAM and MIF estimates for (D) the mean forces and (E, F) the free energies (same color code as above). The error is defined as 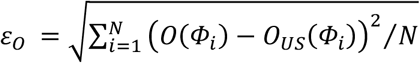, where *O*(*Φ*_*i*_) are the estimates based on FCAM, MIF or the mean biasing potential at discrete *Φ*_*i*_ values, and *O*_*US*_(*Φ*_*i*_) are the corresponding values obtained with US. Lines and shaded regions reflect average error and standard deviations, respectively, when each of 10 Metadynamics trajectories is evaluated individually.

The observation that both FCAM and MIF estimates are rather insensitive to the amplitude of the fluctuations of the biasing potential (**Fig. 3**) probably owes to the simplicity of the test system, which allows for fast equilibration times. In such condition, larger Gaussian heights (up to moderate values, e.g., 0.5 kcal/mol) actually shorten the ‘filling time’ (**Fig. 3F**) and consequently lead to a faster exploration of the space of Φ and more rapid convergence of mean forces and associated free energies (**Fig. 3DE**). A slight accuracy loss is only detected at the largest height value considered (w = 5 kcal/mol, **Fig. 3DE**), for which the time-fluctuations of the biasing potential reach the same order of magnitude of the free-energy changes along Φ, while the average biasing potential still resembles the underlying free energy (**Fig. 3C** and **Fig. S2**). Such deterioration in accuracy is indicative of systematic errors associated with lack of equilibration in the degrees of freedom orthogonal to the biased Φ dihedral angle, and logically is more pronounced for FCAM than MIF, as the latter estimates are not directly dependent on the biasing forces.

An analogous analysis for deca-L-alanine is summarized in **Fig. 4**, based on Metadynamics simulations wherein the end-to-end distance is the biased CV, with Gaussian heights (*w*) ranging from 0.005 to 0.5 kcal/mol (leading to σ_b_ values from 0.5 to 5 kcal/mol (**Fig. S3**)). This analysis confirms that in the regime where the time-fluctuations in the biasing potential are modest, FCAM and MIF estimates converge to the results obtained with US at comparable rates and near identical accuracy (**Fig. 4AB** and Fig. **4DE**). In contrast to what we observed for butyramide, however, both estimates are more sensitive to the magnitude of the time-fluctuations in the biasing potential; accordingly, so are the free-energy estimates based on time-averages of this potential (**Fig. 4CF**). Systematic errors in both FCAM and MIF estimates are apparent for the largest height considered (w = 0.5 kcal/mol), which appear to arise from degrees of freedom orthogonal to the end-to-end distance that cannot reach full equilibration while large time-fluctuations are induced on the biasing potential. More specifically, the mean-force profiles (which are not offset by an arbitrary additive constant) indicate these errors are more pronounced at distances between 16 and 26 Å (**Fig. 4A**); these values correspond to a heterogeneous ensemble of partially extended conformations, which appear to slowly interconvert (not shown). A possible strategy to reduce this type of errors is to expand or optimize the selection of CV so as to include the slowest degrees of freedom, which could be identified, for example, through a spectral gap analysis.^15^ However, as our data shows, a more straightforward solution is simply to limit the time-fluctuations of the biasing potential.

**Figure 4.**
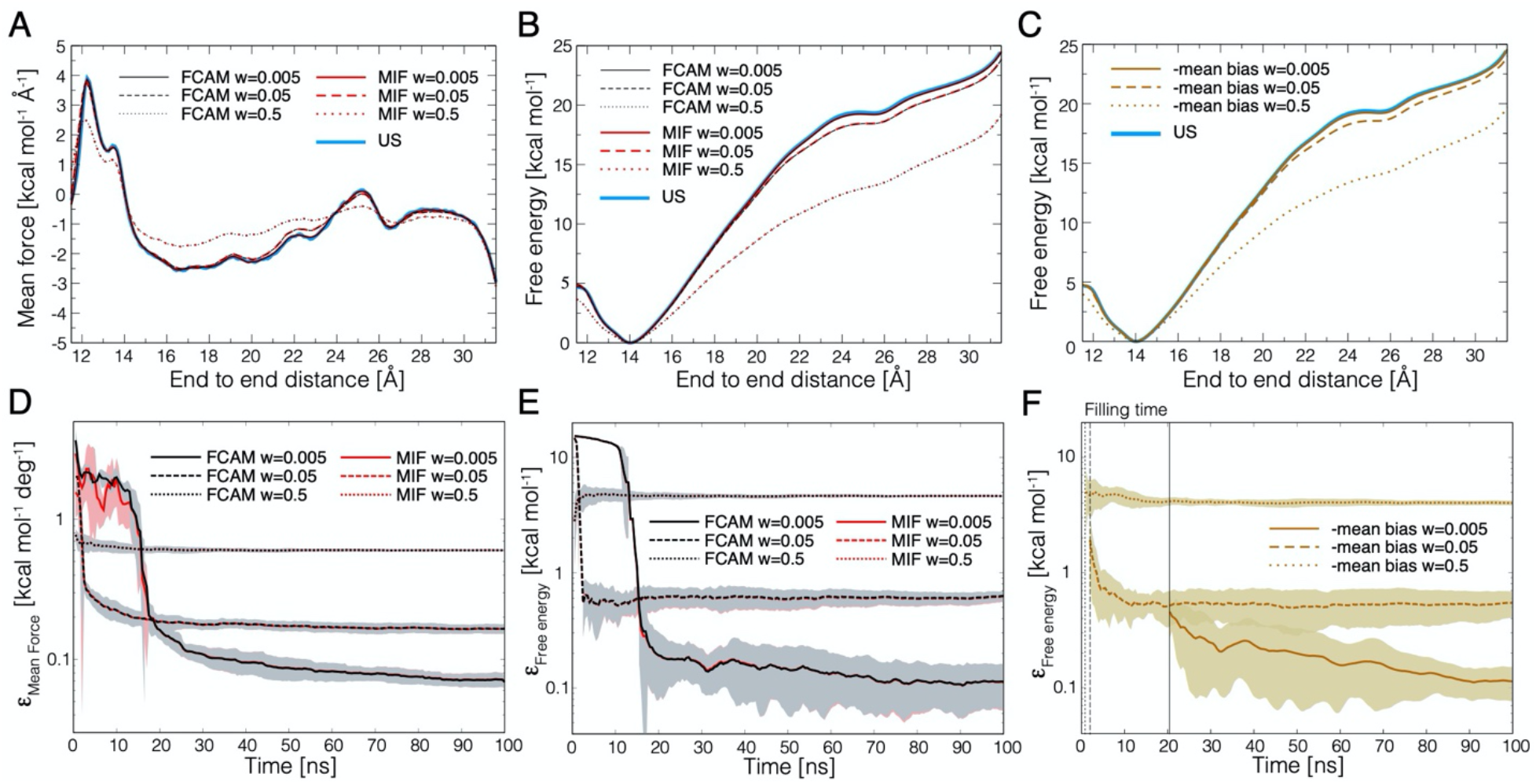
Comparison of FCAM and MIF convergence rates and accuracy, for Metadynamics simulations of deca-L-alanine generated with different amplitudes of the time-fluctuations of the biasing potential acting on the end-to-end distance. (**A**) Mean-force profiles and (**B, C**) free-energy profiles estimated with FCAM (Eq. 9, *black*), with MIF (Eq. 10, *red*) and from the time-average of the Metadynamics biasing potential after the filling time (*brown*), for different values of the height of the Gaussians used to construct that biasing potential (*w*). Each profile was calculated from the combined sampling of 10 independent Metadynamics trajectories of 100 ns each. Analogous profiles obtained with an umbrella-sampling (US) calculation (same as Fig. 2) are shown for comparison (*blue*). (**D, E, F**) Sampling-time dependence of the error in the FCAM and MIF estimates for (D) the mean forces and (E, F) the free energies (same color code as in panels A-C), defined as in Fig. 3. Lines and shaded regions reflect average error and standard deviations, respectively, when each of 10 Metadynamics trajectories is evaluated individually.

### Two-dimensional free energy surfaces from concurrent one-dimensional biases

We next assessed FCAM for cases where multiple one-dimensional biases are implemented on the same trajectory. To this end, we generated 10 independent simulations (of 30 ns each) of solvated butyramide using concurrent Metadynamics biases^21^ that enhance sampling of both Φ and Ψ (**Fig. 1**). That is, two biasing potentials *V*_1_(Φ, *t*) and *V*_2_(Ψ, *t*) were constructed and applied independently, and therefore the overall bias potential is *V*(Φ, Ψ, *t*) = *V*_1_(Φ, *t*) + *V*_2_(Ψ, *t*). This biasing scheme produces a uniform distribution on Φ and Ψ individually, but owing to the correlations between these two CVs, the corresponding two-dimensional space is not sampled uniformly (**Fig. S4**). That is, the overall bias potential, *V*(Φ, Ψ, *t*), does not compensate the free-energy surface along Φ and Ψ, and hence it is not a valid free-energy estimator. Similarly, the biasing potentials *V*_1_(Φ, *t*) and *V*_2_(Ψ, *t*) do not compensate the individual projections of this free-energy surface along either Φ or Ψ. To verify whether the correct two-dimensional free energy can be nevertheless recovered with FCAM, we mapped the space of Φ and Ψ on a regular grid of spacing 2.5° in each direction and for each grid point we calculated the mean force using Eq. 9; for comparison, we also calculated MIF estimates based on the same sampling (using Eq. 10).^55^ In both cases we then obtained a free-energy surface by integration, and assessed its accuracy against the result from an independent US calculation.

In **Fig. 5AB** we report mean-force surfaces for each Φ and Ψ component and the associated free-energy landscape, obtained with FCAM from the combined sampling of the concurrent Metadynamics simulations. The mean-force surfaces are smooth and well-defined throughout, except near the central region, i.e, (Φ, Ψ)~(0,0), which features the highest free-energy barrier and by construction is poorly sampled in the Metadynamics trajectories (**Fig. S4**). Nonetheless, mean-forces calculated using FCAM and MIF generally show an excellent correlation (**Fig. 5C**) and the corresponding free energies match very well the US results (**Fig. 5D**), regardless of the type of estimate used (either FCAM or MIF). Moreover, analysis of the sampling-time dependence of the error in the FCAM and MIF free-energy estimates, with respect to the corresponding US values, underscores both methodologies converge to the expected result with similar rate (**Fig. 5E**). It is thus clear FCAM is a suitable unbiasing methodology for this kind of trajectory data.

**Figure 5.**
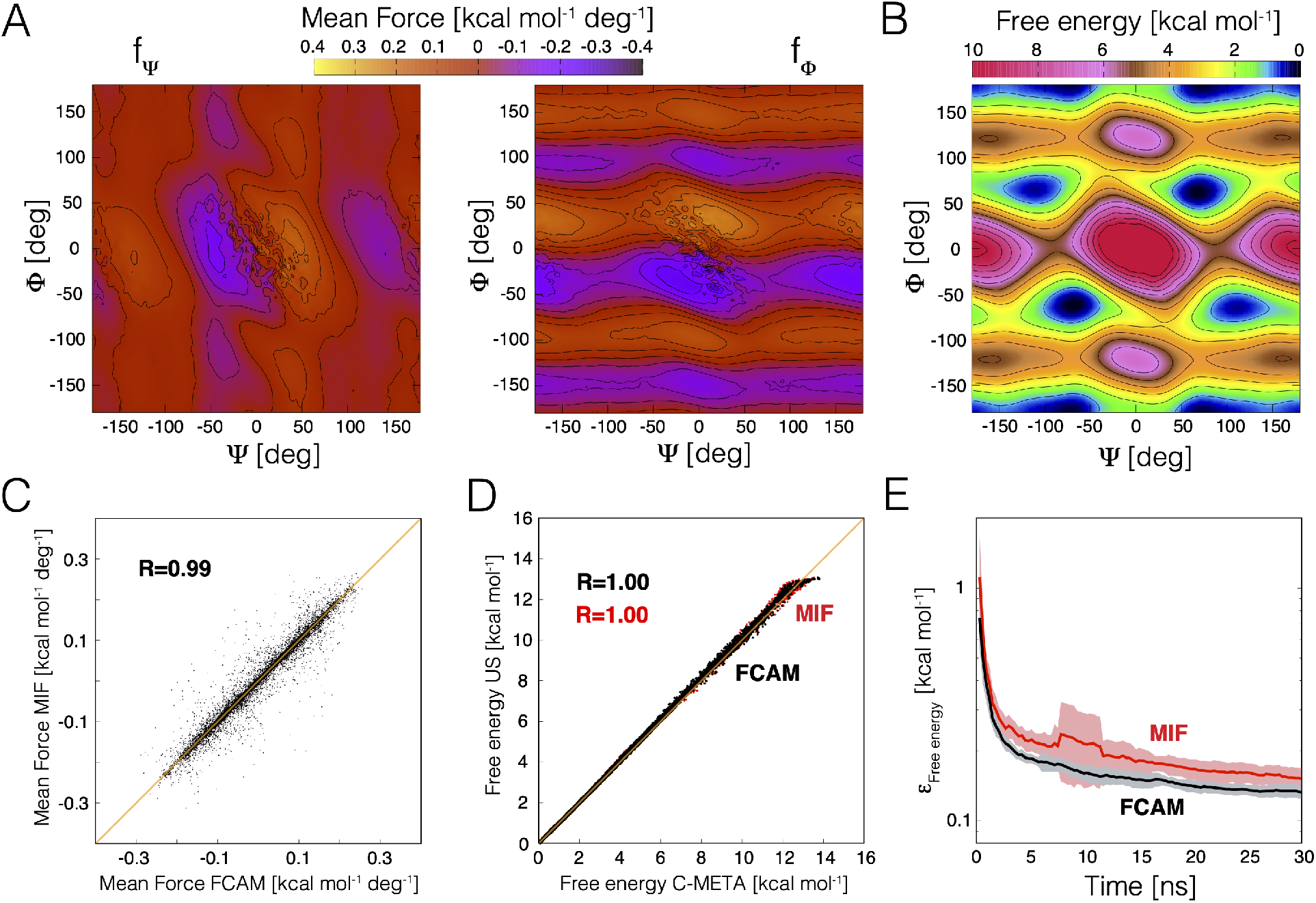
Convergence and accuracy of FCAM for trajectories of solvated butyramide using concurrent Metadynamics biases on the Φ and Ψ dihedral angles. (**A**) Mean-force surface along Φ and Ψ (f_Φ_, f_Ψ_), obtained with FCAM from 10 trajectories of 30 ns each. Contour lines are drawn every 0.05 kcal mol^−1^ deg^−1^. The space of Φ and Ψ was mapped on a lattice of spacing 2.5° in each direction; for each lattice point we calculated the mean force using Eq. 9, using as a density estimator the product of two Gaussians of standard deviation 2.5°. (**B**) Free-energy surface derived from the mean-force surfaces shown in panel (A), using KMC (Methods). Contour lines are drawn every 1 kcal/mol. (**C**) Correlation between mean forces calculated using FCAM and analogous estimates calculated using MIF ^55^ (Eq. 10, and a density estimator identical to that used for FCAM). Data for Φ and Ψ components are combined. (**D**) Correlation between free energies calculated with FCAM or MIF, based on the concurrent Metadynamics trajectories, and analogous data calculated with an US simulation of 130 ns implementing a static biasing potential on Φ and Ψ (constructed using a two-dimensional Metadynamics simulation of 75 ns designed to explore uniformly the Φ and Ψ space). (**E**) Time dependence of the error in the free-energy estimates based on FCAM or MIF, relative to the US free-energy estimates (defined as in Fig. 3). Lines and shaded regions reflect average error and standard deviations when the 10 trajectories are evaluated independently.

### Two-dimensional free-energy surfaces from coupled one-dimensional biases

As mentioned, the ultimate goal of this development is to formulate a robust methodology for computing multidimensional free-energy landscapes from multiple simulations that apply adaptive biasing potentials to different CVs. For example, in Bias-Exchange (BE) Metadynamics^20^ many simulations are carried out in parallel, coupled to one another through a replica-exchange scheme, and each applying Metadynamics biasing potentials to related or entirely different CVs (or to the same CVs but with different Gaussian heights or widths). To date, multidimensional free-energy landscapes based on this type of complex data have been typically derived using WHAM (Eq. 4 and Eq. 5)^25^, but as noted above this approach is somewhat ill-defined in the context of adaptive-biasing methods.

To begin to evaluate the capability of FCAM to extract free-energy landscapes from this kind of simulations, we first considered the solvated alanine dipeptide as a model system. Specifically, we carried out a BE simulation using two coupled replicas (200 ns each); both replicas were biased with standard Metadynamics, but one is enhanced to explore the Φ Ramachandran torsional angle, while the other targets Ψ (**Fig. 6**). We also carried an analogous simulation wherein both replicas implemented a well-tempered Metadynamics bias, which differs from the standard method in that the height of the added Gaussians gradually diminishes as the simulation progresses. ^10, 11, 41^ In both cases, we used FCAM to derive mean-force and free-energy surfaces on the space of Φ and Ψ, as described above. For comparison, we calculated the same 2D free-energy surface with a conventional MD trajectory of ~2 μs and the resulting probability distribution along Φ and Ψ; and with an US simulation of 240 ns implementing a static biasing potential.

**Figure 6.**
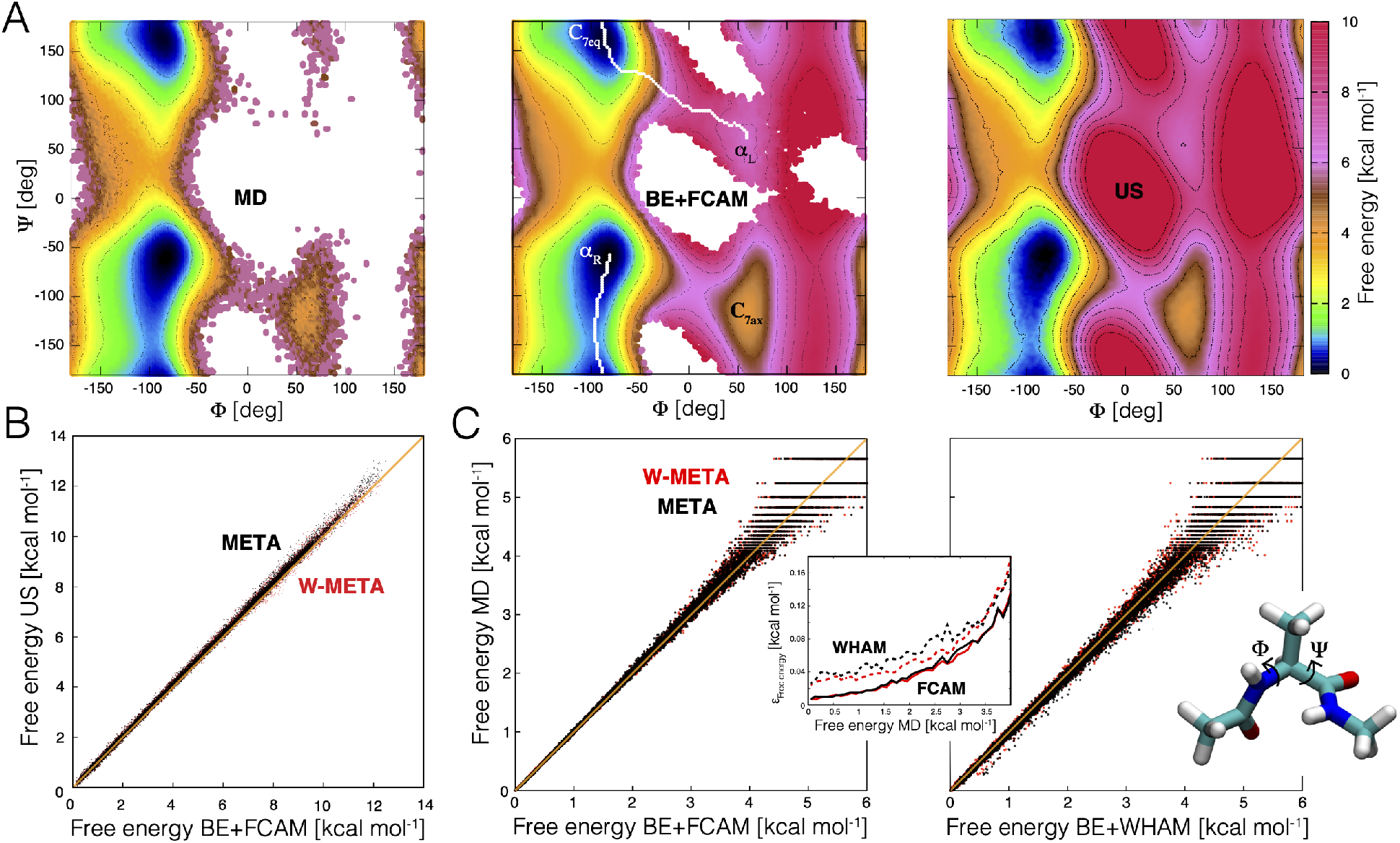
Evaluation of FCAM for adaptively biased simulations of the isomerization of the alanine dipeptide in solvent. (**A**) From left to right, free-energy projections in the space of the Φ and Ψ dihedral angles, calculated with a conventional unbiased MD simulation; with BE Metadynamics simulations, analyzed with FCAM; and with an US simulation implementing a static biasing potential. The free energy is mapped in 2.5° increments in each direction, and contour lines (*black lines*) are drawn at 1 kcal/mol increments. The minimum free energy path (based on Eq.15) connecting α_R_ and α_L_ conformations is indicated on the map calculated with BE/FCAM (*white lines*). The static biasing potential used in the US simulation was constructed from a Metadynamics simulation (of 370 ns) that applies a two-dimensional bias, which explores the *Φ*, Ψ space uniformly. (**B**) Correlation between the free-energy values calculated with FCAM, based on BE Metadynamics simulations, and the corresponding values obtained with US. Data is shown for standard Metadynamics trajectories (META, *black*) and for trajectories obtained with Well-Tempered Metadynamics (W-META, *red*). (**C**) Left, correlation between free-energy values calculated with FCAM, based on BE Metadynamics trajectories (either META or W-META) and the corresponding values calculated from conventional MD. Right, same comparison, based on an alternative derivation of the free energy wherein WHAM is used to unbias the Metadynamics data, rather than FCAM. Inset, absolute values of the deviation in free energy, relative to conventional MD data, in calculations using FCAM (solid line) or WHAM (dashed line), as a function of the magnitude of the free energy. Reported values correspond to a running average over intervals of 0.1 kcal/mol.

As shown in **Fig. 6A**, free-energy landscapes calculated with BE/FCAM and with US closely resemble one another, and that obtained with conventional MD, for the regions that are accessible to each of these methods. By construction, the 2D static biasing potential used in the US simulations leads to sampling of the totality of the Φ–Ψ space, whereas the use of two 1D biasing potentials in the BE Metadynamics simulation precludes the trajectory from reaching the most unfavorable regions. The minimum free-energy path connecting the so-called α_R_ and α_L_ conformations can nevertheless be fully delineated in the BE/FCAM free-energy map (**Fig. 6A**). The excellent correlation between BE/FCAM data and the US calculation is further quantitated in **Fig. 6B**, both for the standard Metadynamics trajectories as well as those obtained with the well-tempered variant. The correlation of the BE/FCAM and conventional MD data is also very good for the range of values where the latter method is reliable (**Fig. 6C**). In this regime, it is also clear that FCAM is systematically more accurate that WHAM, when applied to the same trajectory data (inset of **Fig. 6C**).

### High-dimensional free energy landscapes for both model and complex systems

In the previous sections we have presented evidence for several model systems and demonstrated that FCAM produces accurate free-energy landscapes in one and two dimensions based on simulated trajectories obtained with different adaptive-biasing schemes. For complex systems such as polypeptides and proteins, however, it is rare that only one or two CVs define a mechanism.^1, 3, 69^ Unless these CVs have been specifically optimized so that they collectively capture all the slow degrees of freedom,^15^ a low-dimensional description will often fail to adequately resolve distinct conformational states and the free-energy barriers in between.^10^ A possible solution in such cases is to increase the number of CVs used in the description of the problem (and the bias), but this greater dimensionality also implies a greater computational effort to map the free-energy.^70^ Even for optimized low-dimensional CVs^12–16^, a high-dimensional analysis of the trajectory data is likely to be required to fully characterize the process under study.^22, 47^

To assess whether FCAM may be used to derive accurate free-energy landscapes in multiple dimensions, we applied this method to study the isomerization of solvated Ace-Ala_3_-Nme peptide in a 6-dimensional space.^25^ Specifically, we first carried out a BE Metadynamics simulation using six replicas (260 ns x 6), with each replica adaptively enhanced to sample one of the six Φ and Ψ dihedral angles (**Fig. 7A**). We then used FCAM to analyze these trajectories and derive a free-energy landscape in the space of all six angles. For comparison, we also evaluated this landscape using a conventional unbiased MD simulation. Specifically, we calculated a ~6 μs MD trajectory and derived a six-dimensional probability density based on the same Gaussian density estimator used for computing mean forces (Eq. 1). As shown in **Fig. 7A**, despite the high dimensionality and the angular resolution of our analysis (12°), there is an excellent correlation between the two calculations, especially in the range of free energy values where conventional MD is accurate (< 4 kcal/mol). Free-energy estimates from these two methods differ only by 0.35 kcal/mol on average.

**Figure 7.**
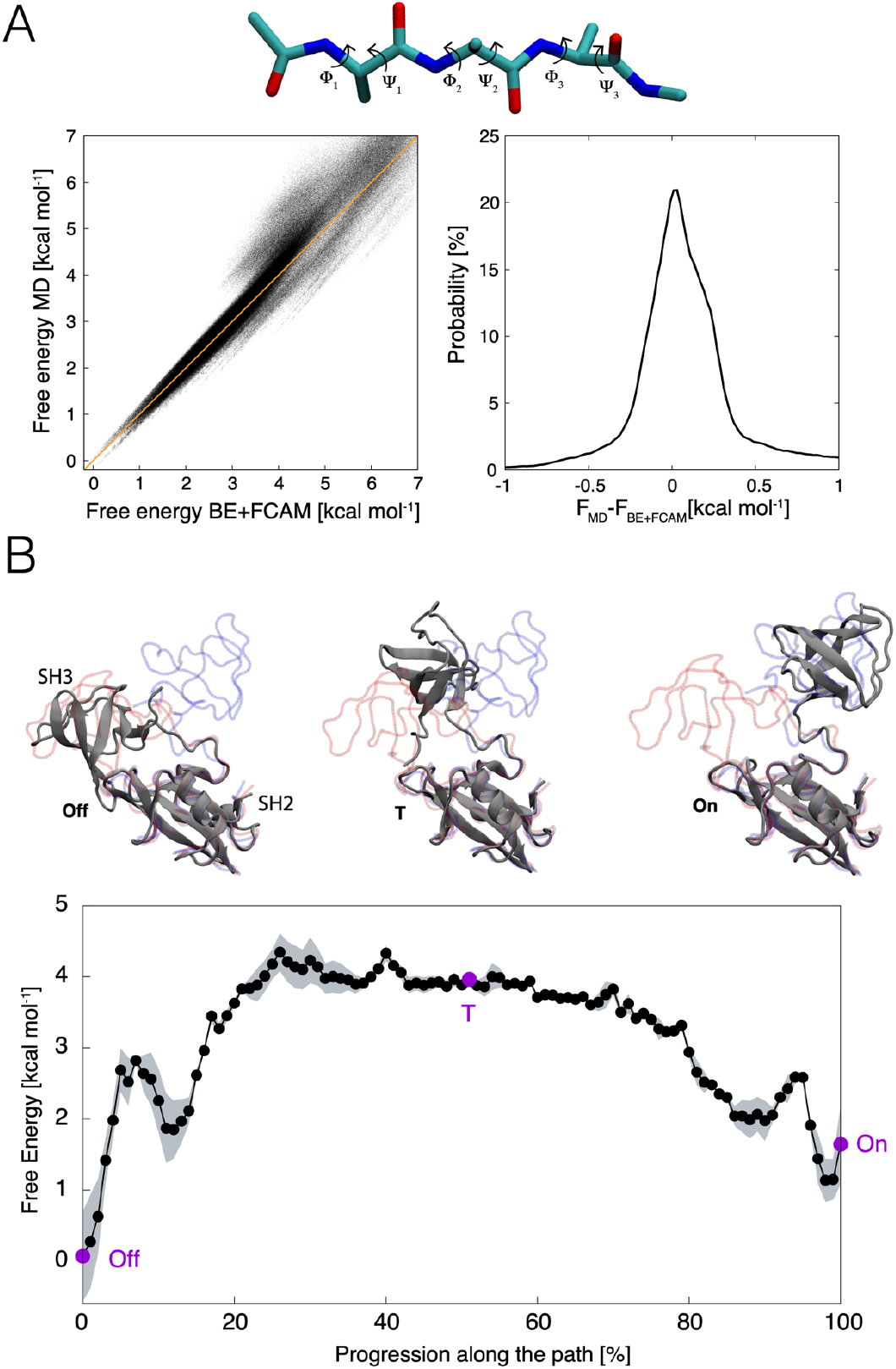
Evaluation of FCAM for adaptively biased simulations in highly dimensional spaces. (**A**) *Top*, Ace-Ala_3_-Nme peptide, highlighting the 6 backbone dihedral angles used as CVs. *Left*, correlation between six-dimensional free-energies values obtained with FCAM, on the basis of BE Metadynamics simulations, and those derived from a conventional MD trajectory of ~6 μs. In both cases, the CV space was mapped with a lattice of 12° spacing in all direction, leading to ~2.5 millions lattice points. The free energy from MD was obtained by mapping a six-dimensional probability density on this lattice, using the same Gaussian density estimator used for FCAM (standard deviation of 12°). Right, histogram of the observed differences (in kcal/mol) between the free-energy estimates based on conventional MD and on BE/FCAM, for the same points in CV space. (**B**) Minimum free-energy path (Eq. 15) connecting the ‘on’ and ‘off’ states of the SH3-SH2 tandem of the Abl kinase, obtained with FCAM on the basis of BE simulations designed to sample the 5-dimensional space.^44^ Selected configurations along the path (purple circles) are shown atop the free-energy plot (gray cartoons). X-ray structures of the ‘on’ and ‘off’ states are overlaid (blue and red, respectively). The shaded area reflects the standard error of the 5-dimensional free-energy profile, obtained through block analysis.

To evaluate FCAM for a more complex biomolecular system, we also examined the conformational landscape of the SH3-SH2 domain-tandem of the Abl tyrosine kinase (**Fig. 7B**), using trajectory data previously reported.^44^ This tandem is known to spontaneously interconvert between two conformers, one inhibitory or “on” and the other non-inhibitory or “off”. The existing simulation data consists of 32 trajectories (of 200 ns each), calculated with BE Metadynamics. This calculation was designed to explore five CVs describing the position and orientation of one domain relative to the other; specifically, each of the replicas was enhanced to sample different combinations these CVs, in pairs ^44^. Using FCAM (Eq. 9 and KMC) we derived the free energy as a function of all five CVs and identified the minimum free energy pathway between the ‘on’ and ‘off’ states (**Fig. 7B**). Consistent with our previous analysis, which had been based on WHAM ^44^, as well as with experimental data,^71^ the results show the isolated tandem favors the off conformation, by 1–1.5 kcal/mol, while the barrier in between the two states is ~4 kcal/mol (**Fig. 7B**). Identical conclusions can be made from evaluation of this transition in a lower-dimensional projection of the free-energy landscape (e.g. 3D instead of 5D), and from examination of alternative and yet significantly probable transition pathways (**Fig. S5**).

## Conclusions

In this work we have introduced a methodology for the calculation of multidimensional free-energy landscapes of complex molecular systems, which we refer to as Force Correction Analysis Method (FCAM). This approach entails a post-hoc derivation of unbiased mean-force estimates from simulation samples obtained with adaptively-biased MD simulations, and builds on concepts outlined in previous studies, such as umbrella integration^32^ and its variants^33–36^. Through a series of applications to both simple and complex systems we have demonstrated that this methodology provides a unified, self-consistent framework to derive thermodynamic quantities from this class of enhanced-sampling techniques. We have also shown that FCAM is compatible with protocols based on either single or multiple trajectories, carried out independently or coupled through an exchange scheme, and implementing biases on the same or different collective variables (CVs). The methodology relies on local averages to calculate mean forces; hence, its practical application is straightforward and only requires quantities that can be deduced readily from trajectory data, or that are already made available by MD simulation programs. Nonetheless, correct application of this analysis method requires that the simulation data be obtained with biases that do not change abruptly or too frequently so that the molecular system remains close to equilibrium. Here, we illustrated the application of the FCAM method for up to six-dimensions. Higher dimensionalities are in principle also amenable, provided that there is enough sampling to perform accurate local averages in connected regions of the CVs space. Specifically tuned estimators of the probability density^47^ might be instrumental in such cases.

For the systems analyzed in this work, the FCAM method showed similar accuracy and convergence rate to an alternative analysis that relies on the estimation of mean instantaneous forces (MIF).^6–8, 53–55^ In contrast to FCAM, MIF estimates do not directly depend on the applied biasing forces but reflect instead the collective effect of the interatomic forces encoded in the simulation energy function. Hence, MIF estimates will typically converge faster than those obtained with FCAM if the biasing potentials are applied in manner that leads to large time-fluctuations. As mentioned, however, this type of fluctuations should be avoided as they drive the molecular system out of equilibrium and lead to sampling errors. By contrast, we have shown that when the biasing scheme is adequate, the convergence and accuracy of FCAM and MIF estimates are on par. Moreover, it is important to note that MIF estimates are practically feasible only for a limited number of CV and simulation specifications. We therefore posit that FCAM offers a more generic and flexible route. In the context of adaptive sampling, FCAM also significantly diminishes the potential for statistical errors, as compared to the WHAM method.^24, 25, 32^

In conclusion, we anticipate that FCAM will greatly facilitate quantitative mechanistic analysis of complex processes in chemical and biological systems. The overall methodology, including the kinetic Monte Carlo scheme used to obtain free energies from the mean forces and to calculate minimum-energy paths, are implemented in python programs freely available through GitHub [https://github.com/Faraldo-Gomez-Lab-at-NIH/Download].

## Acknowledgements

This work was funded by the Division of Intramural Research of the National Heart, Lung and Blood Institute (NHLBI), National Institutes of Health (USA). Computational resources were in part provided by the NIH high-performance computing facility Biowulf.

## References

1. Ma, A.; Dinner, A. R., Automatic method for identifying reaction coordinates in complex systems. Journal of Physical Chemistry B 2005, 109 (14), 6769–6779.

2. Berezhkovskii, A.; Szabo, A., One-dimensional reaction coordinates for diffusive activated rate processes in many dimensions. Journal of Chemical Physics 2005, 122 (1).

3. Best, R. B.; Hummer, G., Reaction coordinates and rates from transition paths. P Natl Acad Sci USA 2005, 102 (19), 6732–6737.

4. Chipot, C.; Pohorille, A., Free Energy Calculations. Springer-Verlag Berlin Heidelberg: 2007.

5. Torrie, G. M.; Valleau, J. P., Non-Physical Sampling Distributions in Monte-Carlo Free-Energy Estimation - Umbrella Sampling. J Comput Phys 1977, 23 (2), 187–199.

6. Darve, E.; Pohorille, A., Calculating free energies using average force. Journal of Chemical Physics 2001, 115 (20), 9169–9183.

7. Henin, J.; Chipot, C., Overcoming free energy barriers using unconstrained molecular dynamics simulations. Journal of Chemical Physics 2004, 121 (7), 2904–2914.

8. Comer, J.; Gumbart, J. C.; Henin, J.; Lelievre, T.; Pohorille, A.; Chipot, C., The adaptive biasing force method: everything you always wanted to know but were afraid to ask. J Phys Chem B 2015, 119 (3), 1129–51.

9. Laio, A.; Parrinello, M., Escaping free-energy minima. Proc Natl Acad Sci U S A 2002, 99 (20), 12562–6.

10. Laio, A.; Gervasio, F. L., Metadynamics: a method to simulate rare events and reconstruct the free energy in biophysics, chemistry and material science. Rep Prog Phys 2008, 71 (12).

11. Bussi, G.; Laio, A., Using metadynamics to explore complex free-energy landscapes. Nat Rev Phys 2020, 2 (4), 200–212.

12. Peters, B.; Trout, B. L., Obtaining reaction coordinates by likelihood maximization. Journal of Chemical Physics 2006, 125 (5).

13. Branduardi, D.; Gervasio, F. L.; Parrinello, M., From A to B in free energy space. J Chem Phys 2007, 126 (5), 054103.

14. Diaz Leines, G.; Ensing, B., Path finding on high-dimensional free energy landscapes. Phys Rev Lett 2012, 109 (2), 020601.

15. Tiwary, P.; Berne, B. J., Spectral gap optimization of order parameters for sampling complex molecular systems. P Natl Acad Sci USA 2016, 113 (11), 2839–2844.

16. Ribeiro, J. M. L.; Bravo, P.; Wang, Y. H.; Tiwary, P., Reweighted autoencoded variational Bayes for enhanced sampling (RAVE). Journal of Chemical Physics 2018, 149 (7).

17. Maragliano, L.; Vanden-Eijnden, E., A temperature accelerated method for sampling free energy and determining reaction pathways in rare events simulations. Chem Phys Lett 2006, 426 (1-3), 168–175.

18. Marinelli, F., Following Easy Slope Paths on a Free Energy Landscape: The Case Study of the Trp-Cage Folding Mechanism. Biophysical Journal 2013, 105 (5), 1236–1247.

19. Pfaendtner, J.; Bonomi, M., Efficient Sampling of High-Dimensional Free-Energy Landscapes with Parallel Bias Metadynamics. Journal of Chemical Theory and Computation 2015, 11 (11), 5062–5067.

20. Piana, S.; Laio, A., A bias-exchange approach to protein folding. J Phys Chem B 2007, 111(17), 4553–9.

21. Gil-Ley, A.; Bussi, G., Enhanced Conformational Sampling Using Replica Exchange with Collective-Variable Tempering (vol 11, pg 1077, 2015). Journal of Chemical Theory and Computation 2015, 11 (11), 5554–5554.

22. Sormani, G.; Rodriguez, A.; Laio, A., Explicit Characterization of the Free-Energy Landscape of a Protein in the Space of All Its Calpha Carbons. J Chem Theory Comput 2020, 16 (1), 80–87.

23. Branduardi, D.; Faraldo-Gomez, J. D., String Method for Calculation of Minimum Free-Energy Paths in Cartesian Space in Freely Tumbling Systems. Journal of Chemical Theory and Computation 2013, 9 (9), 4140–4154.

24. Kumar, S.; Rosenberg, J. M.; Bouzida, D.; Swendsen, R. H.; Kollman, P. A., Multidimensional Free-Energy Calculations Using the Weighted Histogram Analysis Method. J Comput Chem 1995, 16 (11), 1339–1350.

25. Marinelli, F.; Pietrucci, F.; Laio, A.; Piana, S., A kinetic model of trp-cage folding from multiple biased molecular dynamics simulations. PLoS Comput Biol 2009, 5 (8), e1000452.

26. Crespo, Y.; Marinelli, F.; Pietrucci, F.; Laio, A., Metadynamics convergence law in a multidimensional system. Phys Rev E 2010, 81 (5).

27. Branduardi, D.; Bussi, G.; Parrinello, M., Metadynamics with Adaptive Gaussians. Journal of Chemical Theory and Computation 2012, 8 (7), 2247–2254.

28. Rosta, E.; Hummer, G., Free Energies from Dynamic Weighted Histogram Analysis Using Unbiased Markov State Model. Journal of Chemical Theory and Computation 2015, 11 (1), 276–285.

29. Stelzl, L. S.; Kells, A.; Rosta, E.; Hummer, G., Dynamic Histogram Analysis To Determine Free Energies and Rates from Biased Simulations. J Chem Theory Comput 2017, 13 (12), 6328–6342.

30. Wu, H.; Mey, A. S. J. S.; Rosta, E.; Noe, F., Statistically optimal analysis of state-discretized trajectory data from multiple thermodynamic states. Journal of Chemical Physics 2014, 141 (21).

31. Wu, H.; Paul, F.; Wehmeyer, C.; Noe, F., Multiensemble Markov models of molecular thermodynamics and kinetics. P Natl Acad Sci USA 2016, 113 (23), E3221–E3230.

32. Kastner, J.; Thiel, W., Bridging the gap between thermodynamic integration and umbrella sampling provides a novel analysis method: “Umbrella integration”. J Chem Phys 2005, 123 (14), 144104.

33. Zheng, L. Q.; Yang, W., Practically Efficient and Robust Free Energy Calculations: Double-Integration Orthogonal Space Tempering. Journal of Chemical Theory and Computation 2012, 8 (3), 810–823.

34. Fu, H. H.; Shao, X. G.; Chipot, C.; Cai, W. S., Extended Adaptive Biasing Force Algorithm. An On-the-Fly Implementation for Accurate Free-Energy Calculations. Journal of Chemical Theory and Computation 2016, 12 (8), 3506–3513.

35. Lesage, A.; Lelievre, T.; Stoltz, G.; Henin, J., Smoothed Biasing Forces Yield Unbiased Free Energies with the Extended-System Adaptive Biasing Force Method. Journal of Physical Chemistry B 2017, 121 (15), 3676–3685.

36. Marinova, V.; Salvalaglio, M., Time-independent free energies from metadynamics via mean force integration. J Chem Phys 2019, 151 (16), 164115.

37. Maragliano, L.; Fischer, A.; Vanden-Eijnden, E.; Ciccotti, G., String method in collective variables: minimum free energy paths and isocommittor surfaces. J Chem Phys 2006, 125 (2), 24106.

38. Marinelli, F.; Faraldo-Gomez, J. D., Ensemble-Biased Metadynamics: A Molecular Simulation Method to Sample Experimental Distributions. Biophys J 2015, 108 (12), 2779–82.

39. Vaneerden, J.; Briels, W. J.; Harkema, S.; Feil, D., Potential of Mean Force by Thermodynamic Integration - Molecular-Dynamics Simulation of Decomplexation. Chem Phys Lett 1989, 164 (4), 370–376.

40. Maragliano, L.; Vanden-Eijnden, E., Single-sweep methods for free energy calculations. Journal of Chemical Physics 2008, 128 (18).

41. Barducci, A.; Bussi, G.; Parrinello, M., Well-tempered metadynamics: A smoothly converging and tunable free-energy method. Physical Review Letters 2008, 100 (2).

42. Biarnes, X.; Pietrucci, F.; Marinelli, F.; Laio, A., METAGUI. A VMD interface for analyzing metadynamics and molecular dynamics simulations. Comput Phys Commun 2012, 183 (1), 203–211.

43. Marinelli, F.; Kuhlmann, S. I.; Grell, E.; Kunte, H. J.; Ziegler, C.; Faraldo-Gomez, J. D., Evidence for an allosteric mechanism of substrate release from membrane-transporter accessory binding proteins. Proc Natl Acad Sci U S A 2011, 108 (49), E1285–92.

44. Corbi-Verge, C.; Marinelli, F.; Zafra-Ruano, A.; Ruiz-Sanz, J.; Luque, I.; Faraldo-Gomez, J. D., Two-state dynamics of the SH3-SH2 tandem of Abl kinase and the allosteric role of the N-cap. P Natl Acad Sci USA 2013, 110 (36), E3372–E3380.

45. Liao, J.; Marinelli, F.; Lee, C.; Huang, Y.; Faraldo-Gomez, J. D.; Jiang, Y., Mechanism of extracellular ion exchange and binding-site occlusion in a sodium/calcium exchanger. Nat Struct Mol Biol 2016, 23 (6), 590–599.

46. Ono, J.; Nakai, H., Weighted histogram analysis method for multiple short-time metadynamics simulations. Chem Phys Lett 2020, 751.

47. Rodriguez, A.; d’Errico, M.; Facco, E.; Laio, A., Computing the Free Energy without Collective Variables. J Chem Theory Comput 2018, 14 (3), 1206–1215.

48. Bonomi, M.; Branduardi, D.; Bussi, G.; Camilloni, C.; Provasi, D.; Raiteri, P.; Donadio, D.; Marinelli, F.; Pietrucci, F.; Broglia, R. A.; Parrinello, M., PLUMED: A portable plugin for free-energy calculations with molecular dynamics. Comput Phys Commun 2009, 180 (10), 1961–1972.

49. Tribello, G. A.; Bonomi, M.; Branduardi, D.; Camilloni, C.; Bussi, G., PLUMED 2: New feathers for an old bird. Comput Phys Commun 2014, 185 (2), 604–613.

50. Bonomi, M.; Bussi, G.; Camilloni, C.; Tribello, G. A.; Banas, P.; Barducci, A.; Bernetti, M.; Bolhuis, P. G.; Bottaro, S.; Branduardi, D.; Capelli, R.; Carloni, P.; Ceriotti, M.; Cesari, A.; Chen, H. C.; Chen, W.; Colizzi, F.; De, S.; De La Pierre, M.; Donadio, D.; Drobot, V.; Ensing, B.; Ferguson, A. L.; Filizola, M.; Fraser, J. S.; Fu, H. H.; Gasparotto, P.; Gervasio, F. L.; Giberti, F.; Gil-Ley, A.; Giorgino, T.; Heller, G. T.; Hocky, G. M.; Iannuzzi, M.; Invernizzi, M.; Jelfs, K. E.; Jussupow, A.; Kirilin, E.; Laio, A.; Limongelli, V.; Lindorff-Larsen, K.; Lohr, T.; Marinelli, F.; Martin-Samos, L.; Masetti, M.; Meyer, R.; Michaelides, A.; Molteni, C.; Morishita, T.; Nava, M.; Paissoni, C.; Papaleo, E.; Parrinello, M.; Pfaendtner, J.; Piaggi, P.; Piccini, G.; Pietropaolo, A.; Pietrucci, F.; Pipolo, S.; Provasi, D.; Quigley, D.; Raiteri, P.; Raniolo, S.; Rydzewski, J.; Salvalaglio, M.; Sosso, G. C.; Spiwok, V.; Sponer, J.; Swenson, D. W. H.; Tiwary, P.; Valsson, O.; Vendruscolo, M.; Voth, G. A.; White, A., Promoting transparency and reproducibility in enhanced molecular simulations. Nat Methods 2019, 16 (8), 670–673.

51. Fiorin, G.; Klein, M. L.; Henin, J., Using collective variables to drive molecular dynamics simulations. Mol Phys 2013, 111 (22-23), 3345–3362.

52. Sidky, H.; Colon, Y. J.; Helfferich, J.; Sikora, B. J.; Bezik, C.; Chu, W. W.; Giberti, F.; Guo, A. Z.; Jiang, X. K.; Lequieu, J.; Li, J. Y.; Moller, J.; Quevillon, M. J.; Rahimi, M.; Ramezani-Dakhel, H.; Rathee, V. S.; Reid, D. R.; Sevgen, E.; Thapar, V.; Webb, M. A.; Whitmer, J. K.; de Pablo, J. J., SSAGES: Software Suite for Advanced General Ensemble Simulations. Journal of Chemical Physics 2018, 148 (4).

53. Carter, E. A.; Ciccotti, G.; Hynes, J. T.; Kapral, R., Constrained Reaction Coordinate Dynamics for the Simulation of Rare Events. Chem Phys Lett 1989, 156 (5), 472–477.

54. Ciccotti, G.; Kapral, R.; Vanden-Eijnden, E., Blue moon sampling, vectorial reaction coordinates, and unbiased constrained dynamics. Chemphyschem 2005, 6 (9), 1809–1814.

55. Henin, J.; Fiorin, G.; Chipot, C.; Klein, M. L., Exploring Multidimensional Free Energy Landscapes Using Time-Dependent Biases on Collective Variables. Journal of Chemical Theory and Computation 2010, 6 (1), 35–47.

56. Phillips, J. C.; Braun, R.; Wang, W.; Gumbart, J.; Tajkhorshid, E.; Villa, E.; Chipot, C.; Skeel, R. D.; Kale, L.; Schulten, K., Scalable molecular dynamics with NAMD. J Comput Chem 2005, 26 (16), 1781–1802.

57. Phillips, J. C.; Hardy, D. J.; Maia, J. D. C.; Stone, J. E.; Ribeiro, J. V.; Bernardi, R. C.; Buch, R.; Fiorin, G.; Henin, J.; Jiang, W.; McGreevy, R.; Melo, M. C. R.; Radak, B. K.; Skeel, R. D.; Singharoy, A.; Wang, Y.; Roux, B.; Aksimentiev, A.; Luthey-Schulten, Z.; Kale, L. V.; Schulten, K.; Chipot, C.; Tajkhorshid, E., Scalable molecular dynamics on CPU and GPU architectures with NAMD. Journal of Chemical Physics 2020, 153 (4).

58. Hénin, J., Fast and accurate multidimensional free energy integration. arXiv:2107.08511 2021.

59. Harland, B.; Sun, S. X., Path ensembles and path sampling in nonequilibrium stochastic systems. J Chem Phys 2007, 127 (10), 104103.

60. MacKerell, A. D.; Bashford, D.; Bellott, M.; Dunbrack, R. L.; Evanseck, J. D.; Field, M. J.; Fischer, S.; Gao, J.; Guo, H.; Ha, S.; Joseph-McCarthy, D.; Kuchnir, L.; Kuczera, K.; Lau, F. T. K.; Mattos, C.; Michnick, S.; Ngo, T.; Nguyen, D. T.; Prodhom, B.; Reiher, W. E.; Roux, B.; Schlenkrich, M.; Smith, J. C.; Stote, R.; Straub, J.; Watanabe, M.; Wiorkiewicz-Kuczera, J.; Yin, D.; Karplus, M., All-atom empirical potential for molecular modeling and dynamics studies of proteins. Journal of Physical Chemistry B 1998, 102 (18), 3586–3616.

61. Mackerell, A. D.; Feig, M.; Brooks, C. L., Extending the treatment of backbone energetics in protein force fields: Limitations of gas-phase quantum mechanics in reproducing protein conformational distributions in molecular dynamics simulations. J Comput Chem 2004, 25 (11), 1400–1415.

62. Hustedt, E. J.; Marinelli, F.; Stein, R. A.; Faraldo-Gomez, J. D.; McHaourab, H. S., Confidence Analysis of DEER Data and Its Structural Interpretation with Ensemble-Biased Metadynamics. Biophys J 2018, 115 (7), 1200–1216.

63. Hess, B.; Kutzner, C.; van der Spoel, D.; Lindahl, E., GROMACS 4: Algorithms for Highly Efficient, Load-Balanced, and Scalable Molecular Simulation. J Chem Theory Comput 2008, 4 (3), 435–47.

64. Duan, Y.; Wu, C.; Chowdhury, S.; Lee, M. C.; Xiong, G. M.; Zhang, W.; Yang, R.; Cieplak, P.; Luo, R.; Lee, T.; Caldwell, J.; Wang, J. M.; Kollman, P., A point-charge force field for molecular mechanics simulations of proteins based on condensed-phase quantum mechanical calculations. J Comput Chem 2003, 24 (16), 1999–2012.

65. Park, S.; Khalili-Araghi, F.; Tajkhorshid, E.; Schulten, K., Free energy calculation from steered molecular dynamics simulations using Jarzynski’s equality. Journal of Chemical Physics 2003, 119 (6), 3559–3566.

66. Raiteri, P.; Laio, A.; Gervasio, F. L.; Micheletti, C.; Parrinello, M., Efficient reconstruction of complex free energy landscapes by multiple walkers metadynamics. Journal of Physical Chemistry B 2006, 110 (8), 3533–3539.

67. Laio, A.; Rodriguez-Fortea, A.; Gervasio, F. L.; Ceccarelli, M.; Parrinello, M., Assessing the accuracy of metadynamics. Journal of Physical Chemistry B 2005, 109 (14), 6714–6721.

68. Bussi, G.; Laio, A.; Parrinello, M., Equilibrium free energies from nonequilibrium metadynamics. Physical Review Letters 2006, 96 (9).

69. Ren, W.; Vanden-Eijnden, E.; Maragakis, P.; E, W. N., Transition pathways in complex systems: Application of the finite-temperature string method to the alanine dipeptide. Journal of Chemical Physics 2005, 123 (13).

70. Scott, D. W., Multivariate density estimation: theory, practice, and visualization. Wiley & Sons: John Hoboken, NJ, USA, 2015.

71. Fushman, D.; Xu, R.; Cowburn, D., Direct determination of changes of interdomain orientation on ligation: Use of the orientational dependence of N-15 NMR relaxation in Abl SH(32). Biochemistry-Us 1999, 38 (32), 10225–10230.

