## Supplementary Figures for "Force-Correction Analysis Method for Derivation of Multidimensional Free Energy Landscapes from Adaptively Biased Replica Simulations"

### Supporting Information

Fabrizio Marinelli[1]* and José D. Faraldo-Gómez [1]*

[1]Theoretical Molecular Biophysics Laboratory
National Heart, Lung and Blood Institute
National Institutes of Health, Bethesda, MD 20814

*Correspondence should be addressed to:
Fabrizio Marinelli or
José D. Faraldo-Gómez

September 4th, 2021

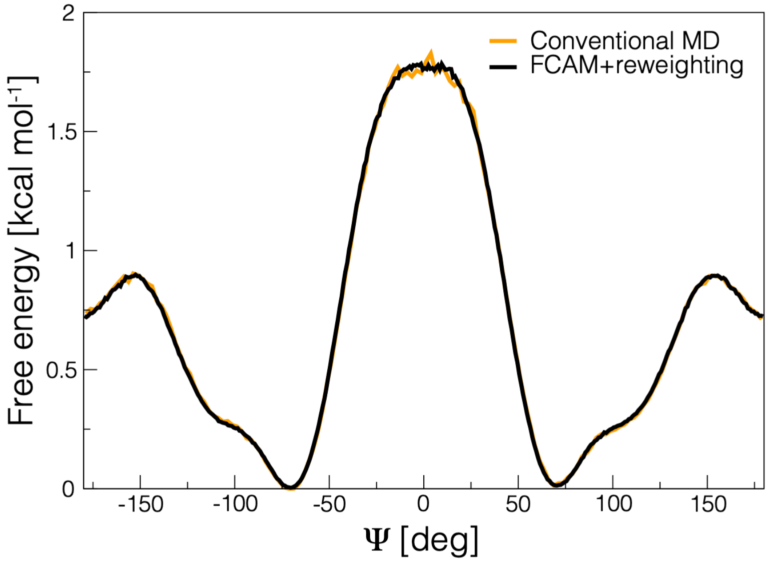


**Figure S1**. Evaluation of the ensemble reweighting scheme (see **Eq. 14)** for Metadynamics simulations of solvated butyramide. The plot compares free-energy profiles along the Ψ dihedral angle, calculated with a conventional 400-ns MD simulation[1] (orange line); and with the FCAM method (black line), based on 10 Metadynamics trajectories (30 ns each) with the Φ dihedral angle as the enhanced CV, followed by ensemble reweighting according to **Eq. 14**.

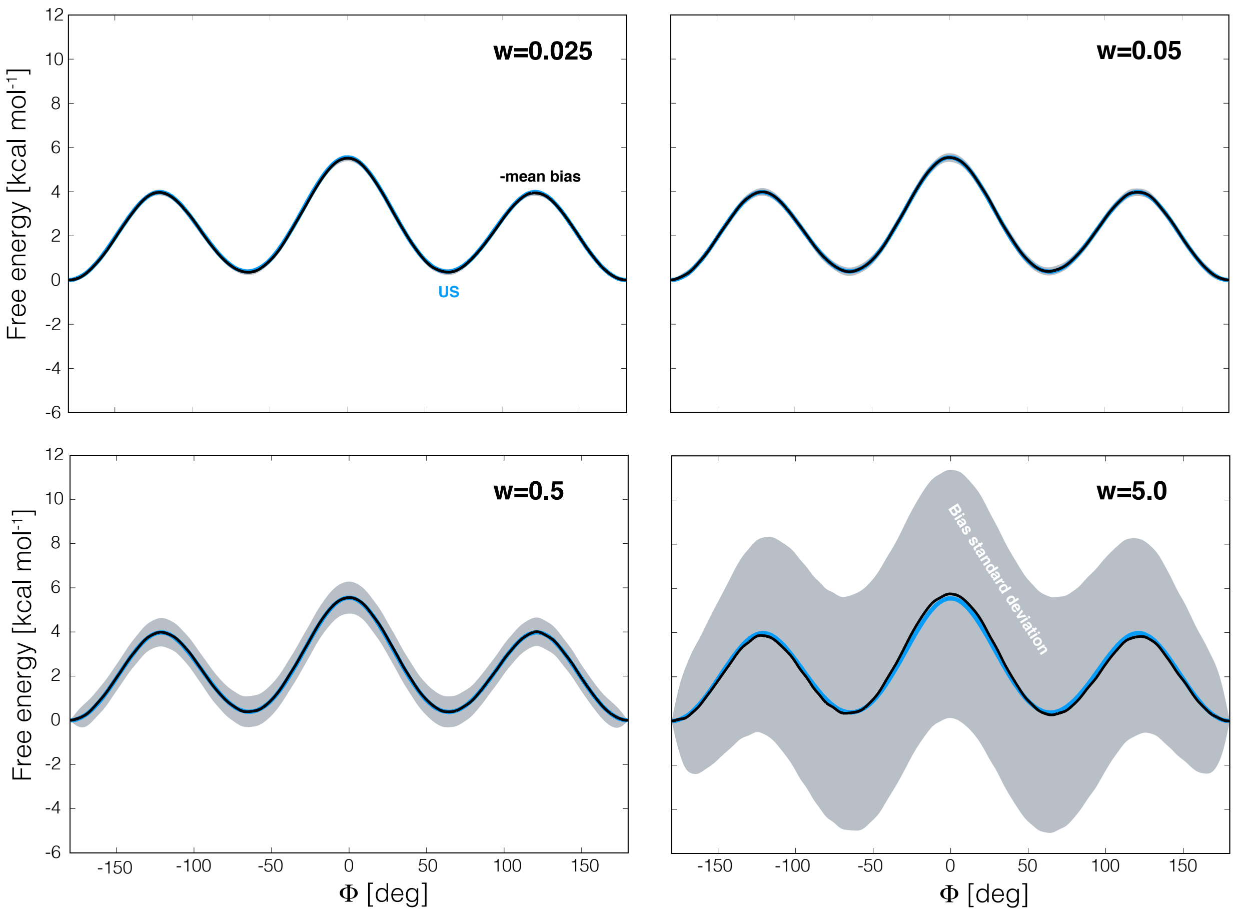


**Figure S2.** Metadynamics simulations of solvated butyramide, with the Φ dihedral angle as the enhanced CV (see **Fig. 1**). The plots quantify the change in amplitude in the time-fluctuations of the biasing potential when Gaussians of increasing height (w) are used to construct this potential. In each plot, the negative of the time-average of the biasing potential after the filling time (*black*) is compared with a free-energy profiles calculated with umbrella sampling (*blue*; same **as Fig. 1** and **Fig. 3**). The gray shaded region indicates in each case the standard deviation of the time-fluctuations of the biasing potential.

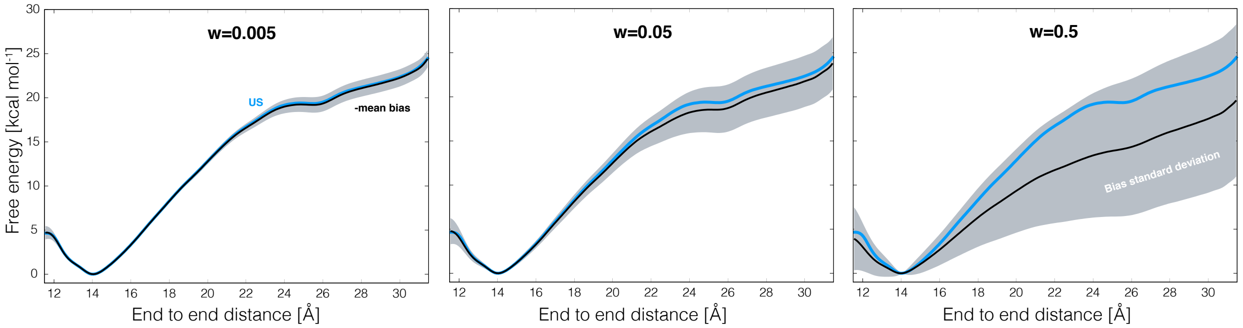


**Figure S3.** Metadynamics simulations of deca-L-alanine, with the end-to-end distance as the enhanced CV (see **Fig. 2**). The plots quantify the change in amplitude in the time-fluctuations of the biasing potential when Gaussians of increasing height (w) are used to construct this potential. In each plot, the negative of the time-average of the biasing potential after the filling time (*black*) is compared with a free-energy profiles calculated with umbrella sampling (*blue*; same as **Fig. 2** and **Fig. 4**). The gray shaded region indicates in each case the standard deviation of the time-fluctuations of the biasing potential.

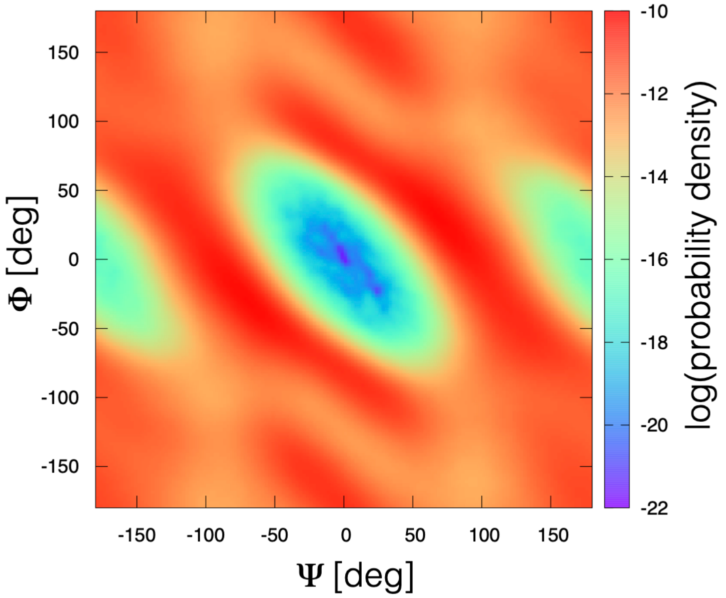


**Figure S4**. Metadynamics simulations of solvated butyramide with concurrent biases applied to the Φ and Ψ dihedral angles (see **Fig. 5**). The plot quantifies the logarithm of the probability density along Φ and Ψ, based on the cumulative sampling obtained with 10 independent trajectories of 30 ns each.

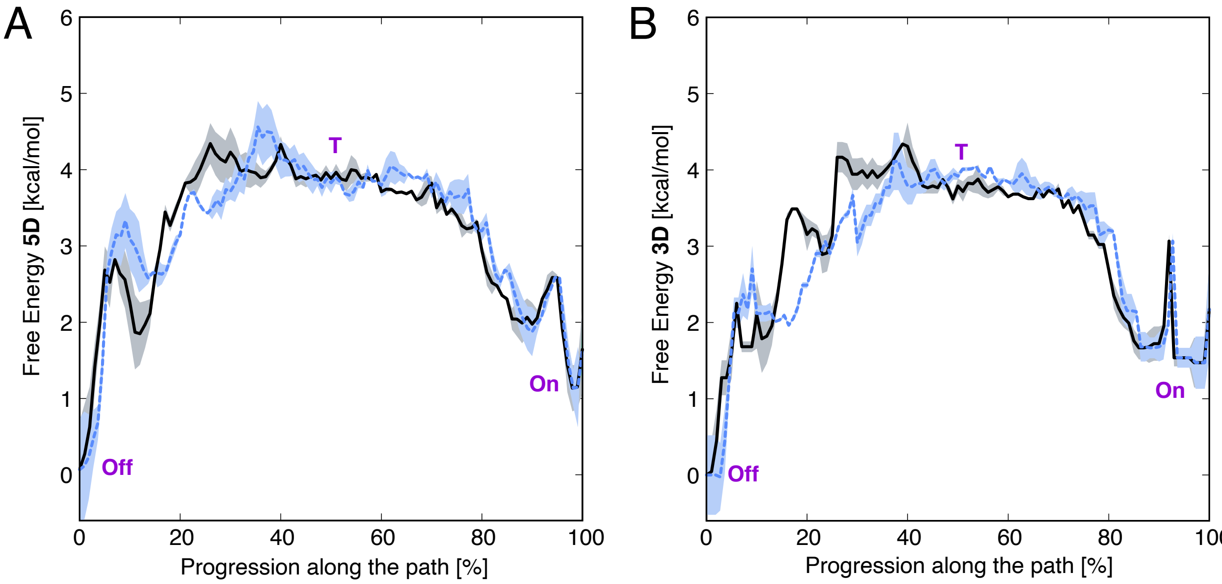


**Figure S5**. Most and second-most probable transition pathways between 'on' and 'off' states of the regulatory SH32 tandem Abl kinase (*black* and *blue-dotted*, respectively), projected in either a (**A**) five- or (**B**) three-dimensional CV space. The underlying free-energy surfaces were obtained through FCAM analysis of published Bias-exchange Metadynamics simulations sampling the five-dimensional space.[2] The two pathways shown in (A) were derived using the kinetic Monte Carlo approach described in Methods, and correspond to pathway probabilities (**Eq. 15**, in log scale) of -451 (*black* line) and -465 (*blue-dotted*). In panel (B**)** the same pathways are mapped onto the three-dimensional free energy landscape obtained by integrating out two of the CV used to define the five-dimensional probability density, specifically the pseudo-dihedral angles used to describe the relative orientation of the SH2 and SH3 orientations.[2] The shaded areas reflect the standard errors based on blocks analysis.

### References


1. Hustedt, E. J.; Marinelli, F.; Stein, R. A.; Faraldo-Gómez, J. D.; McHaourab, H. S., Confidence Analysis of DEER Data and Its Structural Interpretation with Ensemble-Biased Metadynamics. *Biophys J* **2018,** *115*, 1200-1216.
2. Corbi-Verge, C.; Marinelli, F.; Zafra-Ruano, A.; Ruiz-Sanz, J.; Luque, I.; Faraldo-Gómez, J. D., Two-state dynamics of the SH3-SH2 tandem of Abl kinase and the allosteric role of the N-cap. *Proc Natl Acad Sci USA* **2013,** *110*, E3372-E3380.
